# Improved *Aedes aegypti* mosquito reference genome assembly enables biological discovery and vector control

**DOI:** 10.1101/240747

**Authors:** Benjamin J. Matthews, Olga Dudchenko, Sarah Kingan, Sergey Koren, Igor Antoshechkin, Jacob E. Crawford, William J. Glassford, Margaret Herre, Seth N. Redmond, Noah H. Rose, Gareth D. Weedall, Yang Wu, Sanjit S. Batra, Carlos A. Brito-Sierra, Steven D. Buckingham, Corey L Campbell, Saki Chan, Eric Cox, Benjamin R. Evans, Thanyalak Fansiri, Igor Filipović, Albin Fontaine, Andrea Gloria-Soria, Richard Hall, Vinita S. Joardar, Andrew K. Jones, Raissa G.G. Kay, Vamsi K. Kodali, Joyce Lee, Gareth J. Lycett, Sara N. Mitchell, Jill Muehling, Michael R. Murphy, Arina D. Omer, Frederick A. Partridge, Paul Peluso, Aviva Presser Aiden, Vidya Ramasamy, Gordana Rašić, Sourav Roy, Karla Saavedra-Rodriguez, Shruti Sharan, Atashi Sharma, Melissa Laird Smith, Joe Turner, Allison M. Weakley, Zhilei Zhao, Omar S. Akbari, William C. Black, Han Cao, Alistair C. Darby, Catherine Hill, J. Spencer Johnston, Terence D. Murphy, Alexander S. Raikhel, David B. Sattelle, Igor V. Sharakhov, Bradley J. White, Li Zhao, Erez Lieberman Aiden, Richard S. Mann, Louis Lambrechts, Jeffrey R. Powell, Maria V. Sharakhova, Zhijian Tu, Hugh M. Robertson, Carolyn S. McBride, Alex R. Hastie, Jonas Korlach, Daniel E. Neafsey, Adam M. Phllippy, Leslie B. Vosshall

## Abstract

Female *Aedes aegypti* mosquitoes infect hundreds of millions of people each year with dangerous viral pathogens including dengue, yellow fever, Zika, and chikungunya. Progress in understanding the biology of this insect, and developing tools to fight it, has been slowed by the lack of a high-quality genome assembly. Here we combine diverse genome technologies to produce AaegL5, a dramatically improved and annotated assembly, and demonstrate how it accelerates mosquito science and control. We anchored the physical and cytogenetic maps, resolved the size and composition of the elusive sex-determining “M locus”, significantly increased the known members of the glutathione-S-transferase genes important for insecticide resistance, and doubled the number of chemosensory ionotropic receptors that guide mosquitoes to human hosts and egg-laying sites. Using high-resolution QTL and population genomic analyses, we mapped new candidates for dengue vector competence and insecticide resistance. We predict that AaegL5 will catalyse new biological insights and intervention strategies to fight this deadly arboviral vector.

---

Understanding unique aspects of mosquito biology and developing control strategies to reduce their capacity to spread pathogens^1,2^ requires an accurate and complete genome assembly (Fig. 1a). Because the *Ae. aegypti* genome is large (~1.3 Gb) and highly repetitive, the 2007 genome project (AaegL3)^3^ was unable to produce a contiguous genome fully anchored to a physical chromosome map^4^. A more recent assembly, AaegL4^5^, produced chromosome-length scaffolds but suffered from short contigs (contig N50: 84kb) and a correspondingly large number (31,018) of gaps. Taking advantage of the significant advances in sequencing and assembly technology in the decade since the first draft genome of *Ae. aegypti* was published, we used long-read Pacific Biosciences sequencing and Hi-C scaffolding to produce a new reference genome (AaegL5) that is highly contiguous, representing a decrease of 93% in the number of contigs, and anchored end-to-end to the three *Ae. aegypti* chromosomes (Fig. 1, 2a, and Extended Data Fig. 1). Using optical mapping and Linked-Read sequencing, we validated local structure and predicted structural variants between haplotypes. Using this new assembly, we generated a dramatically improved gene set annotation (AaegL5.0), as assessed by a mean increase in RNA-Seq read alignment of 12%, connections between many gene models previously split across multiple contigs, and a roughly two-fold increase in the enrichment of ATAC-Seq alignments near predicted transcription start sites. We demonstrate the utility of AaegL5 and the AaegL5.0 annotation by investigating a number of scientific questions that could not be addressed with the previous genome (Figs 2–5, Extended Data Figs 1–10, Supplementary Data 13–24, and Supplementary Methods and Discussion).

**Figure 1.**
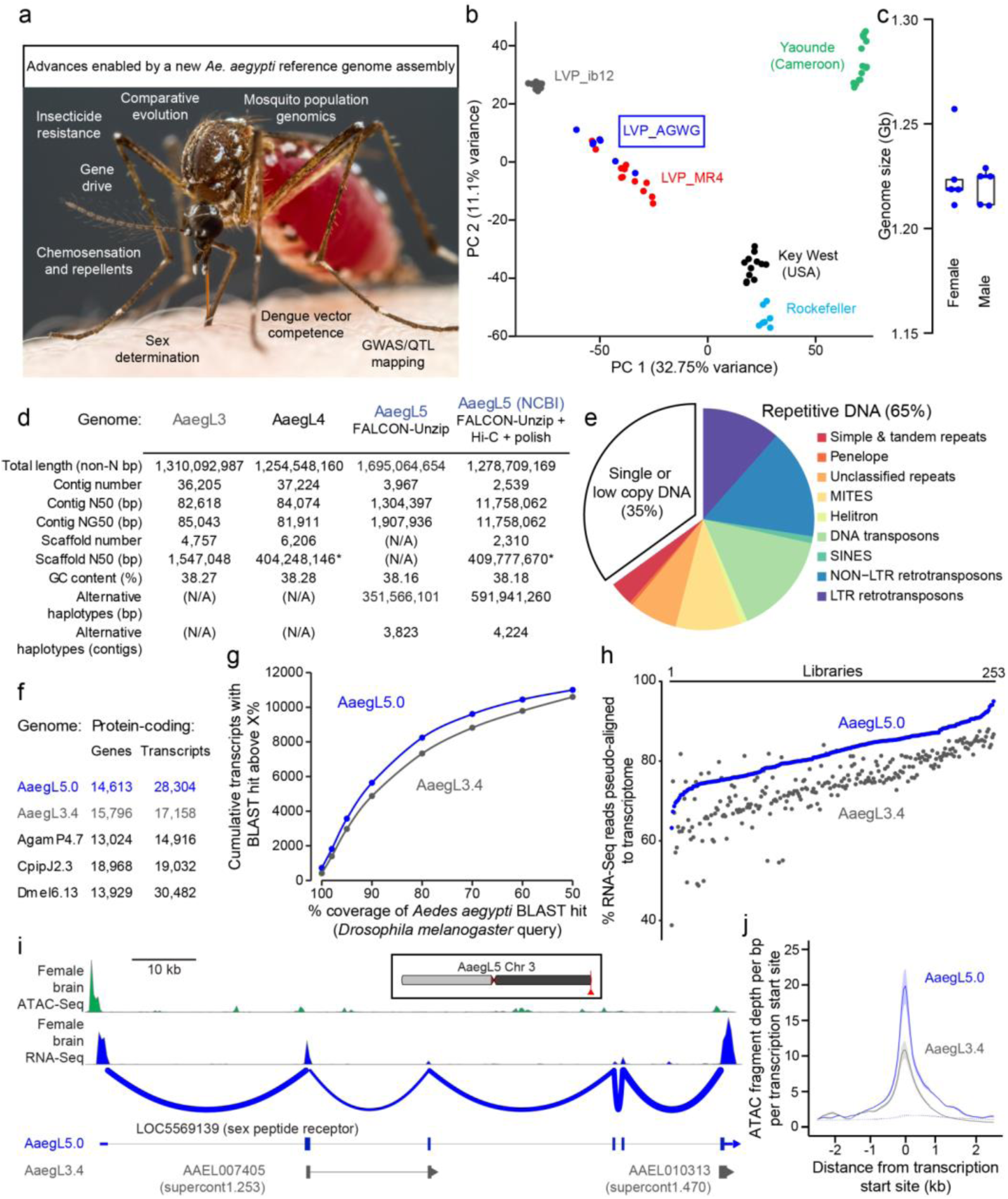
AaegL5 assembly statistics, annotation, and chromatin accessibility analysis. **a**, Visual abstract of utility of the AaegL5 assembly. Photo of a blood-fed *Ae. aegypti* female by Alex Wild. **b**, Principal component analysis (PCA) of allelic variation of the indicated strains at 11,229 SNP loci. **c**, Flow cytometry analysis of LVP_AGWG genome size. Box plot: median= blue line, boxes 1^st^/3^rd^ quartile, whisker 1.5X interquartile interval (Extended Data Fig. 1). **d**, Comparison of assembly statistics (*Scaffold N50 is the length of chromosome 3, N/A: not applicable). **e**, Pie chart of genome composition (Supplementary Data 1–3). **f**, Comparison of protein-coding genes and transcripts in AaegL5.0 (NCBI RefSeq Release 101) and geneset annotations from indicated species. **g**, AaegL3.4 and AaegL5.0 geneset alignment coverage by BLASTp using *D. melanogaster* proteins as queries. **h**, Alignment of 253 RNA-Seq libraries to AaegL3.4 and AaegL5.0 transcriptomes. Each point on the x-axis represents an independent library ordered by increasing alignment to AaegL5.0. (Supplementary Data 4–9). **i**, *SPR* structure in AaegL3.4 and AaegL5.0, and RNA-Seq and ATAC-Seq reads aligned to AaegL5. Blue lines on the RNA-Seq track indicate splice junctions, with the number of reads spanning a junction represented by line thickness. Exons are represented by tall filled boxes and introns by lines. Arrowheads indicate gene orientation. **j**, Average read profiles across promoter regions, defined as the transcription start site ± 2.5 kb. Solid lines represent Tn5-treated native chromatin using the ATAC-Seq protocol (n=4), dotted lines represent Tn5-treated naked genomic DNA (n=1). Shaded regions represent standard deviation.

**Figure 2.**
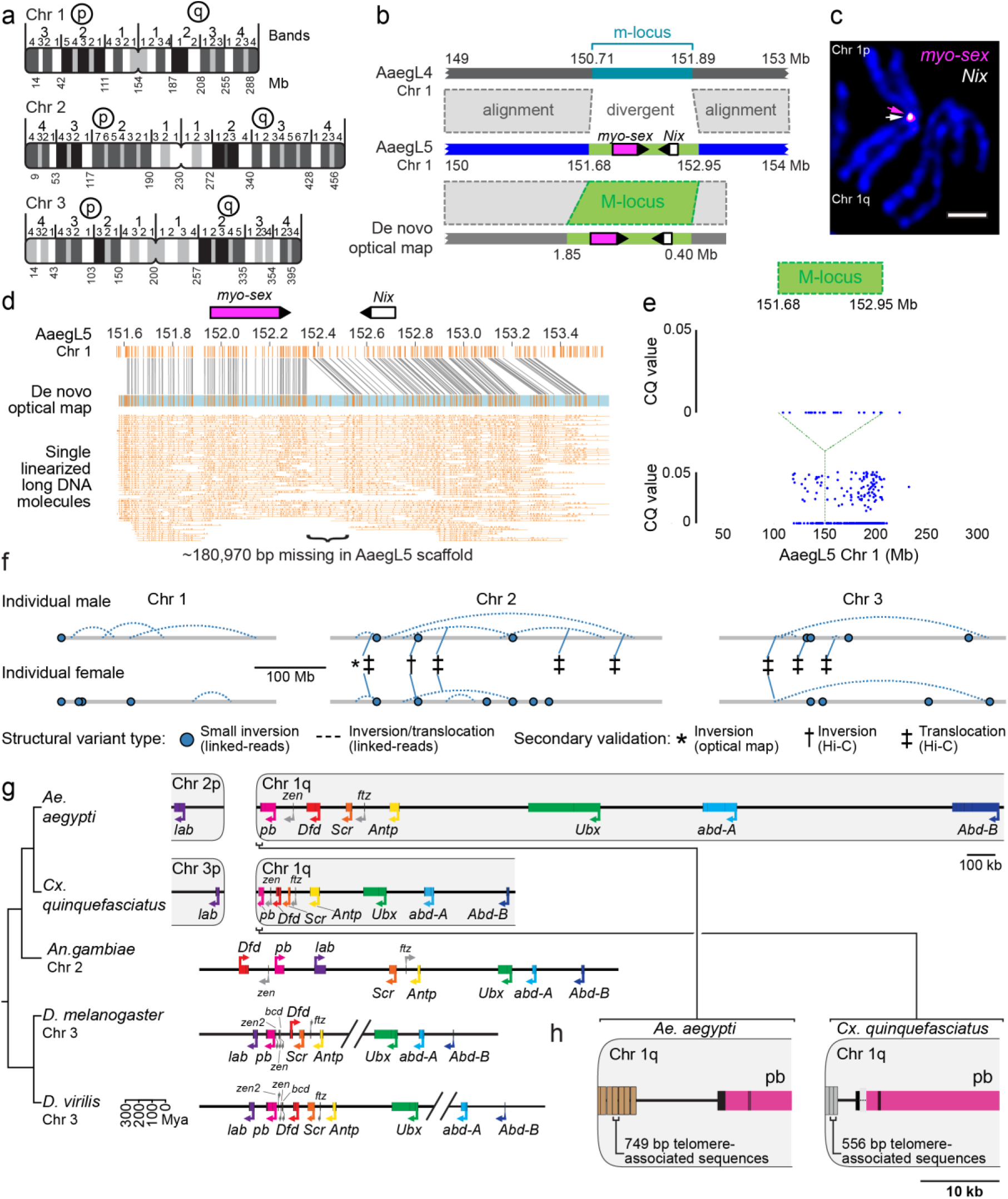
Application of AaegL5 to resolve the sex-determining locus and the HOX gene cluster. **a**, Simplified chromosome map of the *Ae. aegypti* AaegL5 genome assembly. Full map is available in Extended Data Fig. 2 and Supplementary Data 12. **b**, M-locus structure. Grey dashed boxes indicate regions of high identity by alignment. **c**, FISH of BAC clones containing *myo-sex* and *Nix*. Scale bar: 2 µm. **d**, *De novo* optical map spanning the M-locus and bridging the estimated 181 kb gap in the AaegL5 assembly. Single linearized long DNA molecules are cropped at the edges for clarity. **e**, Chromosome-quotient (CQ) analysis of genomic DNA from pure male and female libraries aligned to chromosome 1 of AaegL5. Each dot represents the CQ value of a 1 kb window that was not repeat-masked and had >20 reads aligned from male libraries. **f**, Linked-Reads identified structural variants (SVs) compared to the reference sequence. **g**, Comparative genomic arrangement of the Hox cluster (HOXC) in 5 species (Supplementary Data 14). Due to chromosome arm exchange, Chr 3p in *Cx. quinquefasciatus* is the homologue of Chr 2p in *Ae. aegypti*^5^. **h**, Repeats in putative telomere-associated sequences downstream of *pb* in both species.

**Figure 3.**
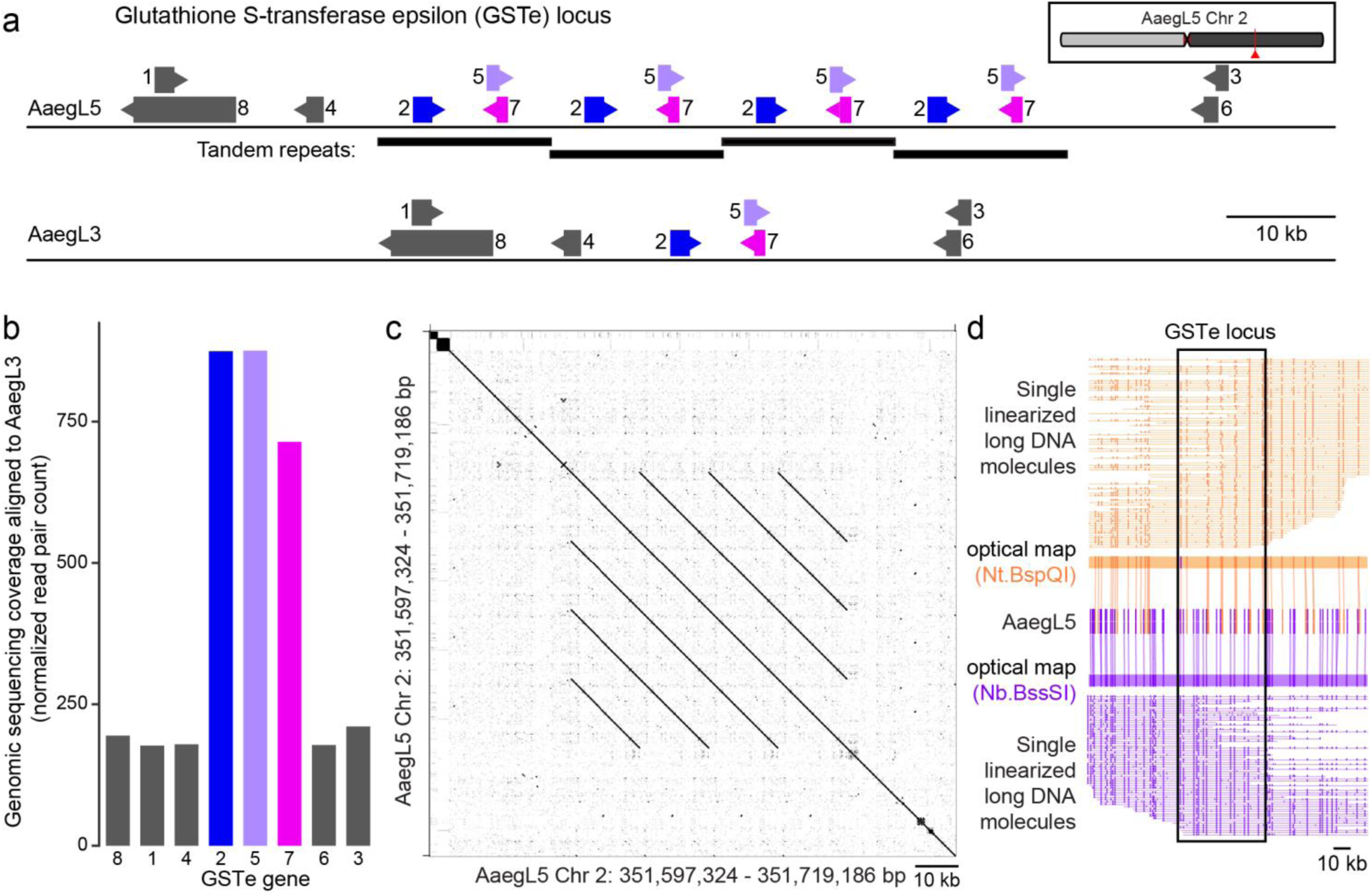
Newly discovered expansion of glutathione S-transferase epsilon gene cluster. **a**, Structure of the glutathione S-transferase epsilon (*GSTe*) gene cluster in AaegL5 compared to AaegL3 (Supplementary Data 15). Arrowheads indicate the direction of transcription for each gene. **b**, Genomic sequencing coverage of *GSTe* genes in AaegL3 (DNA read pairs mapped to each gene, normalized by gene length in kb) from a single LVP_AGWG male. **c**, Alignment of the *GSTe* region of chromosome 2 in AaegL5 to itself, shown as a dot-plot, demonstrates the predicted 4x repeat structure. **d**, Optical mapping of DNA labelled using Nt.BspQI (top) or Nb.BssSI (bottom) provides support for the *GSTe* repeat structure. Individual linearized and labelled long DNA molecules are shown below the map and have been cropped at the edges for clarity.

**Figure 4.**
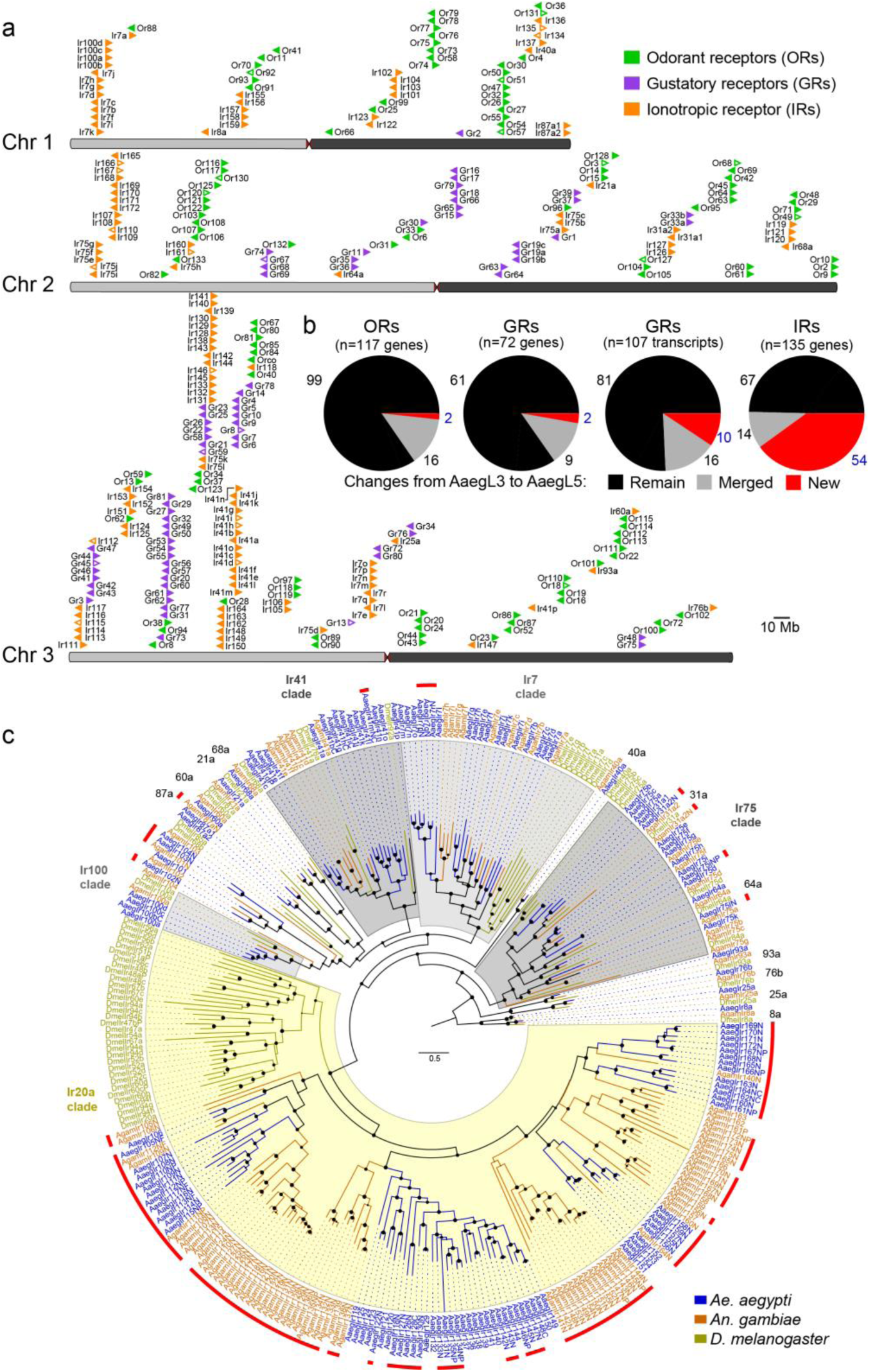
Chromosomal arrangement and expansion in number of chemosensory receptor genes. **a**, The location of predicted chemoreceptors (*ORs*, *GRs*, and *IRs*) across all three chromosomes in AaegL5. The blunt end of each arrowhead marks gene position and arrowhead indicates orientation. Filled and open arrowheads represent intact genes and pseudogenes, respectively (Supplementary Data 20–23). **b**, Pie charts of chemosensory receptor annotation in AaegL3 compared to AaegL5. **c**, Maximum likelihood phylogenetic tree of *IRs* from the indicated species rooted with highly conserved *Ir8a* and *Ir25a* proteins. Conserved proteins with orthologues in all species are named outside the circle, and expansions of *IR* lineages are highlighted with red lines. Suffixes after protein names: C – minor assembly correction, F – major assembly modification, N – new model, and P – pseudogene. Scale bar: amino acid substitutions per site. Filled circles on nodes indicate support levels from approximate likelihood ratio tests from 0–1. Phylogenetic trees of ORs and GRs are in Extended Data Fig.

**Figure 5.**
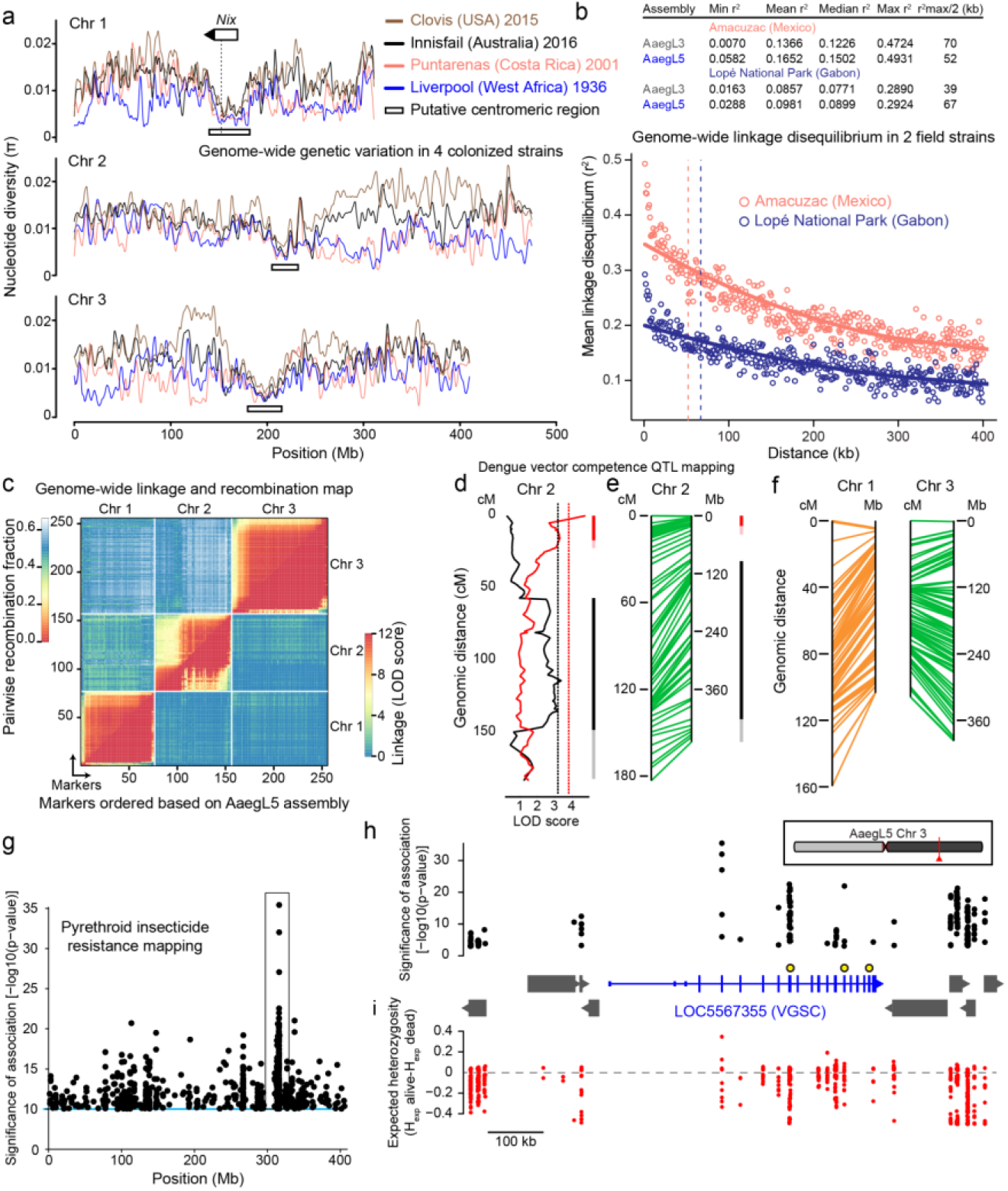
Deploying the AaegL5 genome for applied population genetics. **a**, Chromosomal patterns of nucleotide diversity (π) in four strains of *Ae. aegypti* measured in 100 kb non-overlapping windows and presented as a LOESS-smoothed curve (Extended Data Fig. 9c). **b**, Pairwise linkage disequilibrium (LD) between SNPs located within the same chromosome estimated from 28 wild-caught individuals from the indicated populations. Each point represents the mean LD for that set of binned SNP-pairs. Solid lines are LOESS-smoothed curves, and dashed lines correspond to r ^2 max^ /2 (Extended Data Fig. 9d). **c**, Heat map of linkage based on pairwise recombination fractions for 255 RAD markers ordered according to AaegL5 physical coordinates. **d**, Significant chromosome 2 QTL underlying systemic DENV dissemination in midgut-infected mosquitoes (Extended Data Fig. 10). Curves represent LOD scores obtained by interval mapping and dotted, vertical lines indicate genome-wide statistical significance thresholds (α =0.05). Confidence intervals of significant QTLs (bright colour: 1.5-LOD interval; light colour: 2-LOD interval) are depicted with black representing generalist effects (across DENV serotypes and isolates) and red representing DENV isolate-specific effects (indicative of genotype-by-genotype interactions). **e-f**, Synteny between linkage map (in cM) and physical map (in Mb) for chromosome 2 (**e**) and chromosomes 1 and 3 (**f**). For chromosome 1 there is uncertainty in the cM estimates due to deviations from Mendelian segregation ratios around the M-locus. Because cM and Mb positions are not linearly correlated, confidence intervals in Mb were extrapolated from the cM/Mb synteny of markers delineating the confidence intervals in cM. Number of RAD markers: Chr 1, n=76; Chr 2, n=80; Chr 3, n=99. **g**, SNPs significantly correlated with survival upon exposure to deltamethrin. All SNPs with a –log^10^(prob) > 10 on chromosome 3 are shown. **h**, Zoomed in view of box in (**g**) centred on the newly complete gene model of voltage-gated sodium channel (*VGSC*, transcript variant X3), with SNPs correlated to deltamethrin survival with a –log^10^(prob). Non-coding genes in this interval are omitted for clarity, and all genes but *VGSC* are represented by grey boxes. *VGSC* exons are represented by tall boxes and UTRs by short boxes. Arrowheads indicate gene orientation. The following non-synonymous SNPs in *VGSC* are marked with larger black and yellow circles: (V1016I = 315,983,763, F1534C= 315,939,224, V410L = 316,080,722). Chromosomal position of *VGSC* indicated in red. **i**, Difference in expected heterozygosity (H_exp_ alive – H_exp_ dead) for all SNPs.

We obtained two sources of the laboratory strain LVP_ib12 used for the AaegL3 assembly and found that each differed slightly from the original genome strain (Fig. 1b). We selected one of these and performed 3 rounds of inbreeding, crossing the same male to a single female from three subsequent generations, to generate the LVP_AGWG strain that we used to make Pacific Biosciences sequencing libraries. Animals from the first cross of this inbreeding scheme were used to generate Hi-C, 10X Chromium linked-reads, Illumina paired-end libraries, and Bionano optical maps (Extended Data Fig. 1a). Using flow cytometry, we estimated the genome size of LVP_AGWG as approximately 1.22 Gb (Fig. 1c and Extended Data Fig. 1b). To generate our primary assembly, we produced 166 Gb of Pacific Biosciences data (corresponding to ~130X coverage for a 1.28 Gb genome) and assembled with FALCON-Unzip^6^.

This resulted in a total assembly length of 2.05 Gb (contig N50: 0.96 Mb; NG50: 1.92 Mb). FALCON-Unzip annotated the resulting contigs as either primary (3,967 contigs; N50 1.30 Mb, NG50 1.91 Mb) or haplotigs (3823 contigs; N50 193 kb) representing alternative haplotypes (Fig. 1d and Extended Data Fig. 1e). Notably, the primary assembly was longer than expected for a haploid representation of the *Ae. aegypti* genome as predicted by flow cytometry and prior assemblies. This is consistent with the presence of alternative haplotypes too divergent to be automatically identified as primary/alternative haplotig pairs.

We then combined the primary contigs and haplotigs generated by FALCON-Unzip to create a genome assembly comprising 7790 contigs. We used Hi-C to order and orient these contigs, correct misjoins, and to merge overlaps (Extended Data Fig. 1d, e and Supplementary Methods and Discussion). Briefly, to perform the assembly using Hi-C, we set aside the 359 contigs shorter than 20kb and used Hi-C data to identify 258 misjoins, resulting in 8,306 contigs that we ordered and oriented. Our Hi-C assembly procedure revealed extensive sequence overlap among the contigs, consistent with the assembly of numerous alternative haplotypes. We developed a procedure to merge these alternative haplotypes using Hi-C data, removing 5,440 gaps and boosting the contiguity (NG50: 4.6 Mb; N50, 5.0 Mb). Taken together, the Hi-C assembly procedure placed 94% of sequenced (non-duplicated) bases onto three chromosome-length scaffolds corresponding to the three *Ae. aegypti* chromosomes. After scaffolding, we performed gap-filling and polishing using Pacific Biosciences reads. This removed 270 gaps and further increased the contiguity (NG50: 11.8 Mb; N50, 11.8 Mb), resulting in the final AaegL5 assembly of 1.279 Gb and a complete, gap-free mitochondrial genome. AaegL5 is dramatically more contiguous than the previous AaegL3 and AaegL4 assemblies (Fig. 1d)^3,5^.

Using the TEfam, Repbase, and *de novo* identified repeat databases as queries, we found that 65% of AaegL5 was composed of transposable elements (TEs) and other repetitive sequence (Fig. 1e and Supplementary Data 1–3). Approximately 52% of the assembly is made up of TEs, including ~48% previously identified and ~4% unidentified. The percentage of previously identified TEs is consistent with the 2007 genome, except that P Instability Factor (PIF), a DNA transposable element, increased from 1.1% to 3.3%.

Complete and correct gene models are essential for the study of all aspects of mosquito biology, including the development of transgenic control strategies such as gene drive, while the identification of cis-regulatory elements will aid development of transgenic reagents with cell-type-specific expression. Annotation of AaegL5 was performed using the NCBI RefSeq pipeline and released as annotation version 101 (AaegL5.0; Fig. 1f) followed by manual curation of key gene families. AaegL5.0 formed the basis for a comprehensive quantification of transcript abundance in a series of sex-, tissue-, and developmental stage-specific RNA-Seq libraries (Supplementary Data 4–8). Three lines of evidence indicate that the AaegL5.0 gene-set is substantially more complete and correct than previous versions. First, substantially more genes have high protein coverage when compared to *Drosophila melanogaster* orthologues (915 more genes with at least 80% coverage, a 12.5% increase over AaegL3.4; Fig. 1g). Second, >12% more RNA-Seq reads map to the AaegL5.0 transcriptome than AaegL3.4 (Fig. 1h and Supplementary Data 9). Third, 1463 genes previously annotated separately as paralogues were collapsed into single gene models and 481 previously fragmented gene models were completed by combining multiple partial gene models from the previous assembly due to the increased contiguity of AaegL5 and improvements in annotation methods (Supplementary Data 10 and 11). An example of a now-complete gene model is the sex peptide receptor *(SPR)*, represented by a 6-exon gene model in AaegL5.0 compared to two partial gene fragments on separate scaffolds in AaegL3.4 (Fig. 1i). Splice junctions from RNA-Seq reads fully support the AaegL5.0 gene model and alignments from ATAC-Seq, which are known to co-localise with promoters and other cis-‘/regulatory elements^7^, are consistent with the updated gene model. Genome-wide, a greater proportion of ATAC-Seq reads from adult female brain localised to predicted transcription start sites in AaegL5.0 than AaegL3.4, consistent with the presence of more complete gene models in AaegL5.0 (Fig. 1j). biology, including the development of transgenic control strategies such as gene drive, while the identification of cis-regulatory elements will aid development of transgenic reagents with cell-type-specific expression. Annotation of AaegL5 was performed using the NCBI RefSeq pipeline and released as annotation version 101 (AaegL5.0; Fig. 1f) followed by manual curation of key gene families. AaegL5.0 formed the basis for a comprehensive quantification of transcript abundance in a series of sex-, tissue-, and developmental stage-specific RNA-Seq libraries (Supplementary Data 4–8). Three lines of evidence indicate that the AaegL5.0 gene-set is substantially more complete and correct than previous versions. First, substantially more genes have high protein coverage when compared to *Drosophila melanogaster* orthologues (915 more genes with at least 80% coverage, a 12.5% increase over AaegL3.4; Fig. 1g). Second, >12% more RNA-Seq reads map to the AaegL5.0 transcriptome than AaegL3.4 (Fig. 1h and Supplementary Data 9). Third, 1463 genes previously annotated separately as paralogues were collapsed into single gene models and 481 previously fragmented gene models were completed by combining multiple partial gene models from the previous assembly due to the increased contiguity of AaegL5 and improvements in annotation methods (Supplementary Data 10 and 11). An example of a now-complete gene model is the sex peptide receptor *(SPR)*, represented by a 6-exon gene model in AaegL5.0 compared to two partial gene fragments on separate scaffolds in AaegL3.4 (Fig. 1i). Splice junctions from RNA-Seq reads fully support the AaegL5.0 gene model and alignments from ATAC-Seq, which are known to co-localise with promoters and other cis-‘/regulatory elements, are consistent with the updated gene model. Genome-wide, a greater proportion of ATAC-Seq reads from adult female brain localised to predicted transcription start sites in AaegL5.0 than AaegL3.4, consistent with the presence of more complete gene models in AaegL5.0 (Fig. 1j).

To validate the quality of the new AaegL5 assembly and develop a fine-scale physical genome map for *Ae. aegypti*, we compared the assembly coordinates of 500 BAC clones with physical mapping by fluorescence *in situ* hybridization (FISH). After filtering out repetitive BAC-end sequences and those with ambiguous FISH signals, 377/387 (97.4%) of probes showed concordance between physical mapping and BAC-end alignment. The 10 remaining discordant signals were not supported by 10X or Bionano analysis, and so likely do not reflect misassemblies in AaegL5. We developed a chromosome map for the AaegL5.0 assembly by assigning the coordinates of each outmost BAC clone within a band to the boundaries between bands (Fig. 2a, Extended Data Fig. 2, and Supplementary Data 12). The genome coverage of this physical map is 95.5%, compared to 45% of a previous assembly^8^, and represents the most complete genome map among any mosquito species^9,10^.

## Resolving the structure of the sex-determining M-locus

Sex determination in *Aedes* and *Culex* mosquitoes is governed by a dominant male-determining factor (M-factor) that resides in a male-determining locus (M-locus) on chromosome 1 (ref. ^11–13^). This chromosome is homomorphic between the sexes except for the M/m karyotype. Despite the recent discovery of the M-factor *Nix* in *Ae. aegypti*^14^, the molecular properties of the M-locus remain unknown. In fact, the *Nix* gene was entirely missing in the previous *Ae. aegypti* genome assemblies^3,5^. We first aligned AaegL5 and AaegL4 and identified a region where these two assemblies diverged that contained *Nix* in AaegL5, reasoning that this may represent the divergent M- and m-locus in AaegL5 and AaegL4 respectively (Fig. 2b). A *de novo* optical map assembly spanned the entire putative AaegL5 M-locus and extended beyond its two borders, providing independent evidence for the structure. We estimated the size of the M-locus at approximately 1.5 Mb, including a gap between contigs that is estimated to be ~181 kb based on the optical map (Fig. 2b, d). We tentatively identified the female m-locus as the region in AaegL4 not shared with the M-containing chromosome 1, although we note that the complete phased structure of the M- and m-loci remain to be determined. *Nix* contains a single intron of 100 kb, while *myo-sex*, a gene encoding a myosin heavy chain protein previously shown to be tightly linked to the M-locus^15^, is approximately 300 kb in length compared to 50 kb for its autosomal paralogue. More than 73.7% of the M-locus is repetitive, and interestingly, LTR-retrotransposons comprise 29.9% of the M-locus compared to 11.7% genome-wide. Chromosomal FISH with *Nix*- and *myo-sex*-containing BAC clone probes^16^ showed that these genes co-localise to the 1p pericentromeric region (1p11) in only one homologous copy of chromosome 1, supporting the placement of the M-locus at this position in AaegL5. We note this is contrary to the previously published placement at 1q21 (ref. ^14^) (Fig. 2c). We also investigated the differentiation between the sex chromosomes in the LVP_AGWG strain (Fig. 2e) using a chromosome quotient method to quantify regions of the genome with strictly male-specific signal^17^. A sex-differentiated region in the AGWG strain extends to a ~100 Mb region surrounding the ~1.5 Mb M-locus. This is consistent with the recent analysis of male-female *F*^ST^ in wild population samples and linkage map intercrosses^18^ and could be explained by a large region of reduced recombination that encompasses the centromere and the M-locus^19^. The first description of a fully assembled M-locus in mosquitoes provides exciting opportunities to study the evolution and maintenance of homomorphic sex-determining chromosomes. It was hypothesized that the sex-determining chromosome of *Ae. aegypti* may have remained homomorphic at least since the evolutionary divergence between the *Aedes* and *Culex* genera more than 50 million years ago^20–22^. With the assembled M-locus, we can investigate how these chromosomes have avoided the proposed eventual progression into heteromorphic sex chromosomes^23^.

## Determining the physical arrangement of structural variation and gene families

Structural variation has been associated with capacity to vector pathogens^24^. To use the AaegL5 assembly to investigate the presence of structural variants (SVs) including insertions, deletions, translocations, and inversions present in individual mosquitoes, we produced ‘read cloud’ Illumina sequencing libraries of Linked-Reads with long-range (~80 kb) phasing information from one male and one female mosquito using the 10X Genomics Chromium platform. We used two different SV calling approaches on these data to exploit the long-range phasing, and also investigated a subset of the SV calls by comparison to Hi-C contact maps and FISH performed with BAC clones predicted to lie within the locus of the SV. We observed abundant small-scale insertions/deletions (indels; 26 insertions and 81 deletions called) and inversions/translocations (29 called) in these two individuals (Fig. 2f and Supplementary Data 13). Eight of the inversions/translocations coincided with structural variants seen by Hi-C or FISH, suggesting that those variants are relatively common within this population and can be detected by different methods. Validation of SVs with other data types indicates that this Linked-Read approach, in conjunction with the AaegL5 assembly, is a promising strategy to investigate structural variation systematically within individuals and populations of *Ae. aegypti*.

*Hox* genes encode highly conserved transcription factors important for specifying the identity of segments along the anterior/posterior body axis of all metazoans^25^. In most vertebrates, *Hox* genes are clustered in a co-linear arrangement, while they are often disorganized or split in other animal lineages^26^. Taking advantage of the chromosome-length scaffolds generated here, we studied the structure of the *Hox* cluster (HOXC) in *Ae. aegypti*. All *Hox* genes in closely-related Dipterans are present as a single copy in *Ae. aegypti*, but we identified a split between *labial* (*lab*) and *proboscipedia* (*pb*) that placed *lab* on a separate chromosome (Fig. 2g and Supplementary Data 14). We confirmed the break using chromosome-length scaffolds in AaegL4, which was generated with Hi-C contact maps from a different *Ae. aegypti* strain^5^. We additionally found that a similar split exists in *Culex quinquefasciatus*, suggesting that the break between *lab* and *pb* occurred before these two species diverged. Although a split between *lab* and *pb* is not unprecedented^27^, a unique feature of this split is that both *lab* and *pb* appear to be close to telomeres. Evidence supporting this view is the presence of long tandem repetitive sequences neighbouring *pb* in both *Ae. aegypti* and *Cx. quinquefasciatus*, reminiscent of telomere-associated sequences in species that lack telomerase^28^ (Fig. 2h).

Glutathione-S-transferases (GSTs) are involved in detoxification of compounds including insecticides. They are encoded by a large gene family comprising several classes, two of which, epsilon and delta, are insect-specific^29^. In numerous species, increased GST activity has been associated with resistance to multiple classes of insecticide, including organophosphates, pyrethroids, and the organochlorine DDT^29^ Amplification of detoxification genes is one mechanism by which insects can develop resistance to insecticides^30^. A cluster of GST epsilon genes on chromosome 2 shows evidence of expansion in AaegL5 relative to AaegL3, either due to strain variation or incorrect assembly of AaegL3 and AaegL4. Three genes located centrally in the cluster (*GSTe2*, *GSTe5*, *GSTe7*) are duplicated four times (Fig. 3a-c and Supplementary Data 15). Short Illumina read coverage and optical maps confirmed the copy number and arrangement of these duplications in AaegL5 (Fig. 3b, d). GSTe2 is a highly efficient metaboliser of DDT^31^, and it is interesting to note that the cDNA from three GST genes in the quadruplication (*GSTe2*, *GSTe5*, *GSTe7*) was detected at higher levels in DDT-resistant *Ae. aegypti* mosquitoes from southeast Asia^32^. It will be important to use the AaegL5 assembly to investigate GST gene sequence and copy number in diverse strains that are resistant or susceptible to insecticides.

## Curation of multi-gene families important for immunity, neuromodulation, and sensory perception

Large multi-gene families are notoriously difficult to assemble and correctly annotate because recently duplicated genes typically share high sequence similarity or can be misclassified as alleles of a single gene. We took advantage of the improved AaegL5 genome and AaegL5.0 annotation to curate genes in large multi-gene families encoding proteases, G protein-coupled receptors, and chemosensory receptors.

Serine proteases mediate immune responses^33^ and contribute to blood protein digestion and oocyte maturation in *Anopheles*^34^, while metalloproteases have been linked to vector competence and mosquito-*Plasmodium* interactions^35^. Gene models for over 50% of the 404 annotated serine proteases and metalloproteases in AaegL3.4 were improved in AaegL5.0. We also describe 49 new serine protease/metalloprotease genes that were either not annotated or not identified as proteases in AaegL3.4 (Supplementary Data 16).

G protein-coupled receptors are a large family of membrane receptors that respond to diverse external and internal sensory stimuli. We provide significant corrections to gene models encoding 10 visual opsins and 17 biogenic amine receptors that detect neuromodulators like dopamine and serotonin (Extended Data Fig. 3b and Supplementary Data 17–19). We discovered two isoforms of the *GPRdop1* (X1 and X2) dopamine and *GPRoar1* (X1 and X2) receptors that respectively possess N-terminal and internal regions unique from that predicted for the AaegL3 models. The chromosome-level resolution of the AaegL5 assembly confirms the previously reported *GPRop1–5* cluster on chromosome 3 (ref. ^36^), which has been suggested to be a duplication event associated with adaptation of mosquitoes to new visual environments.

Three large multi-gene families encoding ligand-gated ion channels, the odorant receptors (*ORs*), gustatory receptors (*GRs*), and ionotropic receptors (*IRs*), collectively allow insects to sense a vast array of chemical cues, including carbon dioxide and human body odour that activate and attract female mosquitoes. We identified a total of 117 *ORs*, 72 *GRs* encoding 107 transcripts, and 135 *IRs* in the AaegL5 assembly (Fig. 4a-b and Supplementary Data 20–23). Remarkably, these include 54 new *IRs* (Fig. 4b and Supplementary Data 20) that nearly double the size of the family in *Ae. aegypti*. This discovery prompted us to reannotate *IR* genes in the African malaria mosquito *An. gambiae* where we found a similarly sized group of 64 new IRs. The majority of the new IR genes in both *Ae. aegypti* and *An. gambiae* fall into a large mosquito-specific clade loosely affiliated with the *Ir20a* clade of taste receptors in *D. melanogaster*^37^ (Fig. 4c). Members of this clade in *Ae. aegypti* tend to be expressed in adult forelegs and midlegs of both sexes, or females only, suggesting a potential role in contact chemosensory behaviours such as the evaluation of potential oviposition sites (Extended Data Fig. 5 and 7). The new assembly allowed us to determine the distribution of chemoreceptors across the three chromosomes of *Ae. aegypti* (Fig. 4a), and we found extensive clustering in tandem arrays. Clustering is particularly extreme on chromosome 3p, where a 109 Mb stretch of DNA houses over a third of all these genes (n=111 genes, Fig. 4a). Moreover, although 71 GRs are scattered across chromosomes 2 and 3, only 1 GR (a subunit of the carbon dioxide receptor *AaegGr2*) is found on chromosome 1 (Fig. 4a). We also inferred new phylogenetic trees for each family to investigate the relationship of these receptors in *Ae. aegypti, An. gambiae* and *D. melanogaster* (Fig. 4c and Extended Data Fig. 4) and revised expression estimates for adult male and female neural tissues using deep RNA-Seq^38^ (Extended Data Figs 5–8). The availability of the full repertoire of these receptors will enable studies of mosquito behaviours including response to hosts^39^ and oviposition sites, and will inform the development of novel strategies to disrupt mosquito biting behaviour.

## Genome-wide genetic variation and linkage disequilibrium in wild and colonized *Ae. aegypti*

Measurement of genetic variation within and between populations is key to inferring ongoing and historic evolution in a species^40^. Mapping genetic polymorphism onto a chromosomal genome assembly enables robust inference of demographic history and the identification of regions with low rates of recombination or under selection. To understand the landscape of genomic diversity in *Ae. aegypti*, which has spread from Africa to tropical and subtropical regions around the world in the last century, we performed whole genome resequencing on four laboratory colonies. Broadly, chromosomal patterns of nucleotide diversity should correlate with regional differences in meiotic recombination rates^41^. The Liverpool and Costa Rica colonies maintain extensive diversity despite being colonized more than a decade ago, but show reduced genome-wide diversity on the order of 30–40% relative to the more recently colonized Innisfail and Clovis (Fig. 5a and Extended Data Fig. 9c).

To investigate linkage disequilibrium (LD) in geographically diverse populations of *Ae. aegypti*, we first mapped SNP markers from an Affymetrix chip designed using the previous genome assembly^42^ to positions on AaegL5. We genotyped 28 individuals from two populations from Amacuzac, Mexico and Lopé National Park, Gabon and calculated pairwise LD of SNPs from 1 kb bins both genome-wide and within each chromosome (Fig. 5b and Extended Data Fig. 9d). The maximum LD in the Mexican population is approximately twice that of the Gabon population, and the ability to detect such differences suggests that this approach will be useful in guiding studies of population genomic structure and other characteristics of geographically and phenotypically diverse populations of *Ae. aegypti*.

## Mapping loci for dengue virus vector competence and pyrethroid resistance

To illustrate the value of the chromosome-wide AaegL5 assembly for QTL mapping, we employed restriction site-associated DNA (RAD) markers with unprecedented density to locate QTLs underlying dengue virus (DENV) vector competence. We identified and genotyped RAD markers in the F_2_ progeny of a laboratory cross between wild *Ae. aegypti* founders from Thailand that we previously created to perform QTL mapping using a low-density microsatellite marker map (Extended Data Fig. 10)^43^. A total of 197 F_2_ females in this mapping population had been scored for DENV vector competence against four different DENV isolates (two isolates from serotype 1 and two from serotype 3). The newly developed linkage map included a total of 255 RAD markers (Fig. 5c) with perfect concordance between genetic distances in centiMorgans (cM) and AaegL5 physical coordinates in Mb (Fig. 5c,e-f). In this cross, we detected two significant QTLs on chromosome 2 underlying the likelihood of DENV dissemination from the midgut (i.e., systemic infection), an important component of DENV vector competence^44^. One QTL was associated with a generalist effect across DENV serotypes and isolates, whereas the other was associated with an isolate-specific effect (Fig. 5d). The AaegL5 assembly allowed accurate delineation of the physical coordinates of the QTL boundaries (Fig. 5d-e). Thus, QTL mapping powered by AaegL5 will make it possible to understand the genetic basis of *Ae. aegypti* vector competence for arboviruses.

Pyrethroids are common insecticides used to combat mosquito vectors including *Ae. aegypti*, and resistance to these compounds is a growing problem in much of the world^45^. Understanding the molecular and genetic mechanisms underlying the development of resistance in different mosquito populations is a critical goal in the efforts to combat arboviral pathogens. Many insecticides act on ion channels, and we curated members of the cys-loop ligand-gated ion channel (cysLGIC) superfamily in the new assembly. We found 22 subunit-encoding cysLGICs (Extended Data Fig. 9a and Supplementary Data 24), of which 14 encode putative nicotinic acetylcholine receptor (nAChR) subunits. nAChRs consist of a core group of subunit-encoding genes (α1 to α8 and β1) that are highly conserved between insect species^46^. Each insect analysed so far has at least one divergent subunit that shows low sequence homology to other nAChR subunits^46^; while *D. melanogaster* possesses only one divergent nAChR subunit, *Ae. aegypti* has five. Insect cysLGICs are targets of widely used insecticides in agricultural and veterinary applications, and we found that these compounds impaired the motility of *Ae aegypti* larvae (Extended Data Fig. 9b). cysLGIC-targeting compounds have potential for re-profiling as mosquito larvicides, and the improved annotation presented here provides an invaluable resource for investigating insecticide efficacy.

To demonstrate how a chromosome-scale genome assembly can reveal genetic mechanisms of insecticide resistance, we collected *Ae. aegypti* in Yucatán, Mexico, where pyrethroid-resistant and - susceptible populations co-exist. After phenotyping for survival upon exposure to deltamethrin, we performed a genome-wide population genetic screen for SNPs correlating with resistance to deltamethrin (Fig. 5g and data not shown). This analysis uncovered an association with non-synonymous changes to three amino acid residues of the voltage-gated sodium channel *VGSC*, a known target of pyrethroids (Fig. 5h). The gene model for *VGSC*, a complex locus spanning nearly 500 kb in AaegL5, was incomplete and highly fragmented in AaegL3. SNPs in this region have a lower expected heterozygosity (H_exp_) in the resistant as compared to the susceptible population, supporting the hypothesis that they are part of a selective sweep for the resistance phenotype that surrounds the *VGSC* locus (Fig. 5i). Accurately associating SNPs with phenotypes requires a fully assembled genome, and we expect that AaegL5 will be critical to understanding the evolution of insecticide resistance and other important traits.

## Summary

The high-quality genome assembly and annotation described here will enable major advances in mosquito biology and has already allowed us to carry out a number of experiments that were previously impossible. The highly contiguous AaegL5 genome permitted high-resolution genome-wide analysis of genetic variation and the mapping of loci for DENV vector competence and insecticide resistance. The new appreciation of a large increase in insecticide-detoxifying GSTe genes and a more complete accounting of cysLGICs will catalyse the search for new resistance-breaking insecticides. A doubling in the known number of chemosensory IRs provides opportunities to link odorants and tastants on human skin to mosquito attraction, a key first step in the development of novel mosquito repellents. Sterile Insect Technique and Incompatible Insect Technique show great promise to suppress mosquito populations^47–49^, but these population suppression methods require that only males are released^50^. A strategy that connects a gene for male determination to a gene drive construct has been proposed to effectively bias the population towards males over multiple generations^51^, and improved understanding of M-locus evolution and the function of its genetic content should facilitate genetic control of this important arbovirus vector.

## Acknowledgements

We thank the following members of the mosquito community for their early role in launching the AGWG: Raul Andino, Scott Emrich, Anthony A. James, Mark Kunitomi, Daniel Lawson, Chad Nusbaum, Dave Severson, and Noah Whiteman. Todd Dickinson, Margot Hartley, and Brandon Rice of Dovetail Genomics participated at early stages of this project. Cori Bargmann, David Botstein, Erich Jarvis, and Eric Lander provided early encouragement and facilitation for this project. Nikka Keivanfar, David Jaffe, and Deanna M. Church of 10X Genomics prepared the DNA for SV analysis by S.N.R. and D.E.N. Ying Song of DowDuPont Inc. provided triflumezopyrim for the experiments in Extended Data Fig. 9b. B.J.M. and L.B.V. thank Amy Harmon of the New York Times, and acknowledge generous pro bono contributions of data and analysis from our corporate collaborators.

## Author Contributions

B.J.M. and L.B.V. conceived the study, coordinated data collection and analysis, designed the figures, and wrote the paper with input from all other authors. B.J.M. developed and distributed animals and/or DNA of the LVP_AGWG strain. Detailed author contributions: Pacific Biosciences sample preparation and sequencing: (P.P., M.L.S., J.M.); Assembly: (Sarah. K., R.H., J.K., Sergey K., A.M.P.); Bionano optical mapping: (A.R.H., S.C., J.L., H.C.); Hi-C sample preparation, scaffolding, and deduplication: (O.D., S.S.B. A.D.O., A.P.A., E.L.A); Fig. 1b (B.R.E., A.G.-S., J.R.P); Fig. 1c (J.S.J.); Fig. 1e (L.Z.); Fig. 1f-g (E.C., V.S.J., V.K.K., M.R.M., T.D.M., B.J.M.); Fig. 1h I.A., O.S.A., J.E.C., A.M.W. B.J.W., R.G.G.K., S.N.M., B.J.M.; Fig. 1i-j (M.H., B.J.M.); Fig. 2a (A.S., I.V.S., M.V.S.); Fig. 2b-e (Z.T., M.V.S., I.V.S., A.S., Y.W., J.T., A.C.D., A.R.H., B.J.M.); Fig. 2f (S.N.R., D.E.N.); Fig. 2g (W.J.G., R.S.M., O.D., B.J.M.); Fig. 3. (G.D.W., B.J.M., A.R.H., Sarah K., A.M.P., Sergey K.); Fig. 4 (C.S.M., H.M.R., Z.Z., N.H.R., B.J.M.); Fig. 5a (J.E.C., A.M.W., B.J.W.,R.G.G.K., S.N.M); Fig. 5b (B.R.E., A.G.-S., J.R.P); Fig. 5c-f (A.F., I.F., T.F., G.R., L.L.); Fig. 5g-i (C.L.C, K.S.-R., W.C.B.IV, B.J.M.); Extended Data Fig. 1a (B.J.M.); Extended Data Fig. 1b (J.S.J.); Extended Data Fig. 1c-d (O.D., E.L.A.); Extended Data Fig 1e (Sarah K., J.K., O.D., E.L.A., Sergey K., A.M.P., B.J.M.); Extended Data Fig. 2 (A.S., I.V.S., M.V.S.); Extended Data Fig. 3a (W.J.G., R.S.M.); Extended Data Fig. 3b (C.A.B.-S., S.S., C.H.); Extended Data Fig. 4–8 (C.S.M., H.M.R., Z.Z., N.H.R., B.J.M.); Extended Data Fig. 9a-b (G.J.L., A.K.J., V.R., S.D.B., F.A.P, D.B.S.); Extended Data Fig. 9c (J.E.C., A.M.W. B.J.W., R.G.G.K., S.N.M.); Extended Data Fig. 9d (B.R.E., A.G.-S., J.R.P); Extended Data Fig. 10 (A.F., I.F., T.F, G.R., L.L.); Supplementary Data 1–3 (L.Z.); Supplementary Data 4–9 (I.A., O.S.A., J.E.C., A.M.W. B.J.W., R.G.G.K., S.N.M., B.J.M.); Supplementary Data 10–11 (E.C., V.S.J., V.K.K., M.R.M., T.D.M); Supplementary Data 12 (A.S., I.V.S., M.V.S.); Supplementary Data 13 (S.N.R., D.E.N.); Supplementary Data 14 (W.J.G., R.S.M.); Supplementary Data 15 (G.D.W., B.J.M.); Supplementary Data 16 (S.R., A.S.R.); Supplementary Data 17–19 (C.A.B.-S., S.S., C.H.); Supplementary Data 20–23 (C.S.M., H.M.R., Z.Z., N.H.R., B.J.M.); Supplementary Data 24 (G.J.L., A.K.J., V.R., S.D.B., F.A.P, D.B.S.). The authors declare competing financial interests. P.P., M.L.S., J.M., Sarah. K., R.H., and J.K. are full-time employees at Pacific Biosciences, a company developing single-molecule sequencing technologies.

## Information

Correspondence and requests for materials should be addressed to B.J.M.

## METHODS

### Ethics information

The participation of humans in blood-feeding mosquitoes during routine colony maintenance at The Rockefeller University was approved and monitored by The Rockefeller University Institutional Review Board (IRB protocol LVO-0652). All human subjects gave their written informed consent to participate.

### Mosquito rearing and DNA preparation

*Ae. aegypti* eggs from a strain labelled “LVP_ib12” were supplied by M.V.S. from a colony maintained at Virginia Tech. We performed a single pair cross between a male and female individual to generate material for Hi-C, Bionano optical mapping, flow cytometry, SNP-chip analysis of strain variance, paired-end Illumina sequencing, and 10X Genomics Linked-Reads (Extended Data Fig. 1a). The same single male was crossed to a single female in two additional generations to generate high-molecular weight (HMW) genomic DNA for Pacific Biosciences long-read sequencing and to establish a colony (LVP_AGWG). Rearing was performed as previously described^38^ and all animals were offered a human arm as a blood source.

### SNP analysis of mosquito strains

Data were generated as described^42^, and PCA analysis using LEA 2.0 available for R v3.4.0 (ref. ^52,53^). The following strains were used: *Ae. aegypti* LVP_AGWG (Samples from the laboratory strain used for the AaegL5 genome assembly, reared as described in Extended Fig. 1a by a single pair mating in 2016 from a strain labelled LVP_ib12 maintained at Virginia Tech); *Ae. aegypti* LVP_ib12 (Laboratory strain, LVP_ib12, provided in 2013 by David Severson, University of Notre Dame), *Ae. aegypti* LVP_MR4 (laboratory strain labelled LVP_ib12 obtained in 2016 from MR4 at the Centers for Disease Control via BEI Resources catalogue # MRA-735), *Ae. aegypti* Yaounde, Cameroon (field specimens collected in 2014 and provided by Basile Kamgang), *Ae. aegypti* Rockefeller (laboratory strain provided in 2016 by George Dimopoulos, Johns Hopkins Bloomberg School of Public Health), *Ae. aegypti* Key West, Florida (field specimens collected in 2016 and provided by Walter Tabachnick).

### Flow cytometry

Genome size was estimated by flow cytometry as described^54^ except that the propidium iodide was added at a concentration of 25 µL/mg, not 50 µL/mg, and samples were stained in the cold and dark for 24 hr to allow the stain to fully saturate the sample. In brief, nuclei were isolated by placing a single frozen head of an adult sample along with a single frozen head of an adult *Drosophila virilis* female standard from a strain with 1C = 328 Mb into 1 ml of Galbraith buffer (4.26 g MgCl_2_, 8.84 g sodium citrate, 4.2 g 3-[*N*-morpholino] propane sulfonic acid (“MOPS”), 1 ml Triton X-100, and 1 mg boiled ribonuclease A in 1 litre of ddH_2_O, adjusted to pH 7.2 with HCl and filtered through a 0.22 μm filter)^55^ and grinding with 15 strokes of the A pestle at a rate of 3 strokes/2 sec. The resultant ground mixture was filtered through a 60 μm nylon filter (Spectrum Labs, CA). Samples were stained with 25 μg of propidium iodide and held in the cold (4°C) and dark for 24 hr at which time the relative red fluorescence of the 2C nuclei of the standard and sample were determined using a Beckman Coulter CytoFlex flow cytometer with excitation at 488 nm. At least 2000 nuclei were scored under each 2C peak and all scored peaks had a CV of 2.5 or less^54,55^. Average channel numbers for sample and standard 2C peaks were scored using CytExpert software version 1.2.8.0 supplied with the CytoFlex flow cytometer. Significant differences among strains were determined using Proc GLM in SAS with both a Tukey and a Sheffé option. Significance levels were the same with either option. Genome size was determined as the ratio of the mean channel number of the 2C sample peak divided by the mean channel number of the 2C *D. virilis* standard peak times 328 Mb, where 328 Mb is the amount of DNA in a gamete of the standard. The following species/strains were used: *Ae. mascarensis* (collected by A. Bheecarry on Mauritius in December 2014. Colonized and maintained by J.R.P.), *Ae. aegypti* Ho Chi Minh City F13 (provided by Duane J. Gubler, Duke-National University of Singapore as F1 eggs from females collected in Ho Chi Minh City in Vietnam, between August and September 2013. Colonized and maintained for 13 generations by A.G.-S.), *Ae. aegypti* Rockefeller (laboratory strain provided by Dave Severson, Notre Dame), *Ae. aegypti* LVP_AGWG: Reared as described in Extended Data Fig. 1a from a strain labelled LVP_ib12 maintained by M.V.S. at Virginia Tech), *Ae. aegypti* New Orleans F8 (collected by D. Wesson in New Orleans 2014. Colonized and maintained by J.R.P. through 8 generations of single pair mating), *Ae. aegypti* Uganda 49-ib-G5 (derived by C.S.M. through 5 generations of full-sibling mating of the U49 colony established from eggs collected by John-Paul Mutebi in Entebbe, Uganda in March 2015).

### Pacific Biosciences library construction, sequencing, and assembly

#### HMW DNA extraction for Pacific Biosciences sequencing

HMW DNA extraction for Pacific Biosciences sequencing was performed using the Qiagen MagAttract Kit (#67563) following the manufacturer’s protocol with approximately 80 male sibling pupae in batches of 25 mg.

#### SMRTbell Library Construction and Sequencing

Three libraries were constructed using the SMRTbell Template Prep Kit 1.0 (Pacific Biosciences). Briefly, genomic DNA (gDNA) was mechanically sheared to 60 kb using the Megaruptor system (Diagenode) followed by DNA damage repair and DNA end repair. Universal blunt hairpin adapters were then ligated onto the gDNA molecules after which non-SMRTbell molecules were removed with exonuclease. Pulse field gels were run to assess the quality of the SMRTbell libraries. Two libraries were size selected using SageELF (Sage Science) at 30 kb and 20 kb, the third library was size selected at 20 kb using BluePippin (Sage Science). Prior to sequencing, another DNA damage repair step was performed and quality was assessed with pulse field gel electrophoresis. A total of 177 SMRT cells were run on the RS II using P6-C4 chemistry and 6 hr movies.

#### Contig Assembly and Polishing

A diploid contig assembly was carried out using FALCON v.0.4.0 followed by the FALCON-Unzip module (revision 74eefabdcc4849a8ceF24d1a1bbb27d953247bd7)^6^. The resulting assembly contains primary contigs, a partially-phased haploid representation of the genome, and haplotigs, which represent phased alternative alleles for a subset of the genome. Two rounds of contig polishing were performed. For the first round, as part of the FALCON-Unzip pipeline, primary contigs and secondary haplotigs were polished using haplotype-phased reads and the Quiver consensus caller^56^. For the second round of polishing we used the “resequencing” pipeline in SMRT Link v.3.1, with primary contigs and haplotigs concatenated into a single reference. Resequencing maps all raw reads to the combined assembly reference with BLASR (v. 3.1.0)^57^, followed by consensus calling with arrow.

### Hi-C sample preparation and analysis

#### Library Preparation

Briefly, insect tissue was crosslinked and then lysed with nuclei permeabilised but still intact. Nuclei were then extracted, and libraries were prepared using a modified version of the *in situ* Hi-C strategy that we optimized for insect tissue^58^. Separate libraries were prepared for samples derived from three individual male pupae. The resulting libraries were sequenced to yield 118M, 249M and 114M reads (coverage: 120X), and processed using Juicer^59^.

#### Hi-C Approach

Using the results of FALCON-Unzip as input, we used Hi-C to correct misjoins, to order and orient contigs, and to merge overlaps (Extended Data Fig. 1e). The Hi-C based assembly procedure we employed is described in the Supplementary Methods and Discussion. Notably, the input assembly we used included for this process removed the distinction between primary contigs and the alternative haplotigs generated by FALCON-Unzip. This was essential because Hi-C data identified genomic loci where the corresponding sequence was absent in the primary FALCON-Unzip contigs, and present only in the haplotigs; the loci would have led to gaps, instead of contiguous sequence, if the haplotigs were excluded from the Hi-C assembly process (Extended Data Fig. 1e).

#### Hi-C Scaffolding

We set aside 359 FALCON-Unzip contigs shorter than 20kb, because such contigs are more difficult to accurately assemble using Hi-C. To generate chromosome-length scaffolds, we used the Hi-C maps and the remaining contigs as inputs to the previously described algorithms^5^. Note that both primary contigs and haplotigs were used as input. We performed quality control, polishing, and validation of the scaffolding results using Assembly Tools (Dudchenko et al., in preparation). This produced 3 chromosome-length scaffolds. Notably, the contig N50 decreased slightly, to 929,392 bp, because of the splitting of misjoined contigs.

#### Hi-C Alternative Haplotype Merging

Examination of the initial chromosome-length scaffolds using Assembly Tools (Dudchenko et al., in preparation) revealed that extensive undercollapsed heterozygosity was present. In fact, most genomic intervals were repeated, with variations, on two or more unmerged contigs. This suggested that the levels of undercollapsed heterozygosity were unusually high, and that the true genome length was far shorter than either the total length of the Pacific Biosciences contigs (2,047 Mb), or the initial chromosome-length scaffolds (1,970 Mb). Possible factors that could have contributed to the unusually high rate of undercollapsed heterozygosity seen in the FALCON-Unzip Pacific Biosciences contigs relative to prior contig sets for *Ae. aegypti* generated using Sanger sequencing (AaegL3)^3^, include high heterozygosity levels in the species and incomplete inbreeding in the samples we sequenced. The merge algorithm described in Dudchenko et al.^5^ detects and merges draft contigs that overlap one another due to undercollapsed heterozygosity. Since undercollapsed heterozygosity does not affect most loci in a typical draft assembly, the default parameters are relatively stringent. We therefore ran the merge algorithm using more permissive merge parameters, but found that the results would occasionally merge contigs that did not overlap. To avoid these false positives, we created a system to manually identify and ‘whitelist’ regions of the genome containing no overlap, based on both Hi-C maps and LASTZ alignments (Extended Data Fig. 1c and Supplementary Methods and Discussion). We then reran the merge step, using the whitelist as an additional input. Finally, we performed quality control of the results using Assembly Tools (Dudchenko et al., in preparation), which confirmed the absence of the undercollapsed heterozygosity that we had previously observed. The resulting assembly contained 3 chromosome-length scaffolds (311 Mb, 474 Mb, and 410 Mb), which spanned 94% of the merged sequence length. Notably, the merging of overlapping contigs using the above procedure frequently eliminated gaps, and thus greatly increased the contig N50, from 929,392 to 4,997,917 bp. The assembly contains three chromosome-length scaffolds and 2,364 small scaffolds, which spanned the remaining 6%. These lengths were consistent with the results of flow cytometry and the lengths obtained in prior assemblies.

### Final gap-filling and polishing

#### Scaffolded Assembly Polishing

Following scaffolding and de-duplication, we performed a final round of arrow polishing. PBJelly^60^ from PBSuite version 15.8.24 was used for gapfilling of the de-duplicated HiC assembly (see Protocol.xml in Supplementary Methods and Discussion). After PBJelly, the liftover file was used to translate the renamed scaffolds to their original identifiers. For this final polishing step (run with SMRT Link v3.1 resequencing), the reference sequence included the scaffolded, gap-filled reference, as well as all contigs and contig fragments not included in the final scaffolds (https://github.com/skingan/AaegL5_FinalPolish). This reduces the likelihood that reads map to the wrong haplotype, by providing both haplotypes as targets for read mapping. For submission to NCBI, two scaffolds identified as mitochondrial in origin were removed (see below), and all remaining gaps on scaffolds were standardized to a length of 100 Ns to indicate a gap of unknown size. The assembly QV was estimated using independent Illumina sequencing data from a single individual male pupa (library H2NJHADXY_1/2). Reads were aligned with bwa mem 0.7.12-r1039 (ref. ^61^). Freebayes v1.1.0–50-g61527c5-dirty (ref. ^62^) was used to call SNPs with the parameters -C 2 -0 -O -q 20 -z 0.10 -E 0 -X -u -p 2 -F 0.6. Any SNP showing heterozygosity (e.g. 0/1 genotype) was excluded. The QV was estimated at 34.75 using the PHRED formula with SNPs as the numerator (597,798) and number of bases with at least 3-fold coverage as the denominator, including alternate alleles (1,782,885,792).

#### Identification of mitochondrial contigs

During the submission process for this genome, two contigs were identified as mitochondrial in origin and were removed from the genomic assembly, manually circularized, and submitted separately. The mitochondrial genome is available as GenBank accession number MF194022.1, RefSeq accession number NC_035159.1.

### Bionano optical mapping

#### High-molecular weight DNA extraction

High-molecular weight (HMW) DNA extraction was performed using the Bionano Animal Tissue DNA Isolation Kit (RE-013–10), with a few protocol modifications. A single-cell suspension was made as follows. 47 mg of frozen male pupae were fixed in 2% v/v formaldehyde in kit Homogenization Buffer (Bionano #20278), for 2 min on ice. The pupae were roughly homogenized by blending for 2 sec, using a rotor-stator tissue homogenizer (TissueRuptor, Qiagen #9001271). After another 2 min fixation, the tissue was finely homogenized by running the rotor-stator for 10 sec. Homogenate was filtered with a 100 µm nylon filter, fixed with ethanol for 30 min on ice, spun down, and washed with more Homogenization Buffer (to remove residual formaldehyde). The final pellet was resuspended in Homogenization Buffer.A single agarose plug was made using the resuspended cells, using the CHEF Mammalian Genomic DNA Plug Kit (BioRad #170–3591), following the manufacturer’s instructions. The plug was incubated with Lysis Buffer (Bionano #20270) and Puregene Proteinase K (Qiagen #1588920) overnight at 50°C, then again the following morning for 2 hr (using new buffer and Proteinase K). The plug was washed, melted, and solubilized with GELase (Epicentre #G09200). The purified DNA was subjected to four hr of drop dialysis (Millipore, #VCWP04700) and quantified using the Quant-iT PicoGreen dsDNA Assay Kit (Invitrogen/Molecular Probes #P11496).

#### DNA labelling

DNA was labelled according to commercial protocols using the DNA Labelling Kit –NLRS (RE-012–10, Bionano Genomics, Inc). Specifically, 300 ng of purified genomic DNA was nicked with 7 U nicking endonuclease Nt.BspQI (New England BioLabs, NEB) at 37°C for 2 hr in NEBuffer3. The nicked DNA was labelled with a fluorescent-dUTP nucleotide analogue using Taq polymerase (NEB) for 1 hr at 72°C. After labelling, the nicks were ligated with Taq ligase (NEB) in the presence of dNTPs. The backbone of fluorescently labelled DNA was counterstained with YOYO-1 (Invitrogen).

#### Data collection

The DNA was loaded onto the nanochannel array of Bionano Genomics IrysChip by electrophoresis of DNA. Linearized DNA molecules were then imaged automatically followed by repeated cycles of DNA loading using the Bionano Genomics Irys system.

The DNA molecules backbones (YOYO-1 stained) and locations of fluorescent labels along each molecule were detected using the in-house software package, IrysView. The set of label locations of each DNA molecule defines an individual single-molecule map.

After filtering data using normal parameters (molecule reads with length greater than 150 kb, a minimum of 8 labels, and standard filters for label and backbone signals), a total of 299 Gb and 259 Gb of data were collected from Nt.BspQI and Nb.BssSI samples, respectively.

#### *De novo* genome map assembly

*De novo* assembly was performed with non-haplotype aware settings (optArguments_nonhaplotype_noES_irys.xml) and pre-release version of Bionano Solve3.1 (Pipeline version 6703 and RefAligner version 6851). Based on Overlap-Layout-Consensus paradigm, pairwise comparison of all DNA molecules was done to create a layout overlap graph, which was then used to create the initial consensus genome maps. By realigning molecules to the genome maps (RefineB P-value = 10e^-11^) and by using only the best match molecules, a refinement step was done to refine the label positions on the genome maps and to remove chimeric joins. Next, during an extension step, the software aligned molecules to genome maps (Extension P-value = 10e^-11^), and extended the maps based on the molecules aligning past the map ends. Overlapping genome maps were then merged using a Merge P-value cutoff of 10e^-15^. These extension and merge steps were repeated five times before a final refinement was applied to “finish” all genome maps (Refine Final P-value = 10e^-11^). Two genome map *de novo* assemblies, one with nickase Nt.BspQI and the other with nickase Nb.BssSI, were constructed. Alignments between the constructed *de novo* genome assemblies and the L5 assembly were done using a dynamic programming approach with a scoring function and a P-Value cutoff of 10e^-12^.

### Transposable element identification

#### Identification of known transposon elements (TE)

We first identified known TEs using RepeatMasker (version 3.2.6)^63^ against the mosquito TEfam (https://tefam.biochem.vt.edu/tefam/, data downloaded on July 2017), a manually curated mosquito TE database. We then ran RepeatMasker using the TEfam database and Repbase TE library (version 10.05). The RepeatMasker was set to default parameters with the –s (slow search) and NCBI/RMblast program (2.2.28).

#### *De novo* repeat family identification

We searched for repeat families and consensus sequences using the *de novo* repeat prediction tool RepeatModeler (version 1.0.8)^64^ using default parameters with RECON (version 1.07) and RepeatScout (1.0.5) as core programs. Consensus sequences were generated and classified for each repeat family. Then RepeatMasker was run on the genome sequences, using the RepeatModeler consensus sequence as the library.

#### Tandem repeats

We also predicted tandem repeats in the whole genome and in the repeatmasked genome using Tandem Repeat Finder (TRF)^65^. Long Tandem copies were identified using the “Match=2, Mismatch=7, Delta=7, PM=80, PI=10, Minscore=50 MaxPeriod=500” parameters. Simple repeats, satellites, and low complexity repeats were found using “Match=2, Mismatch=7, Delta=7, PM=80, PI=10, Minscore=50, and MaxPeriod=12” parameters.

### Generation of RefSeq geneset annotation

The AaegL5 assembly was deposited at NCBI in June of 2017 and annotated using the NCBI RefSeq Eukaryotic gene annotation pipeline^66^. Evidence to support the gene predictions came from over 9 billion Illumina RNA-seq reads, 67k PacBio IsoSeq reads, 300k ESTs, and well-supported proteins from *D. melanogaster* and other insects. Annotation Release 101 was made public in July 2017, and specific gene families were subjected to manual annotation and curation. See also: http://www.ncbi.nlm.nih.gov/genome/annotation_euk/Aedes_aegypti/101/

### Alignment of transcriptome data to AaegL5 and quantification of gene expression

RNA-Seq reads from developmental transcriptome^67^, neurotranscriptome^38^ and tissue-specific libraries produced by Verily Life Sciences were mapped to the RefSeq assembly GCF_002204515.2_AaegL5.0 with STAR aligner (v2.5.3a) (https://doi.org/10.1093/bioinformatics/bts635) using the 2-pass approach. Reads were first aligned in the absence of gene annotations using the following parameters: --outFilterType BySJout; --alignIntronMax 1000000; --alignMatesGapMax 1000000; --outFilterMismatchNmax 999; --outFilterMismatchNoverReadLmax 0.04; --clip3pNbases 1; --outSAMstrandField intronMotif; --outSAMattrIHstart 0; --outFilterMultimapNmax 20; --outSAMattributes NH HI AS NM MD; --outSAMattrRGline; --outSAMtype BAM SortedByCoordinate. Splice junctions identified during the 1^st^ pass mapping of individual libraries were combined and supplied to STAR using the –sjdbFileChrStartEnd option for the second pass. Reads mapping to gene models defined by the NCBI annotation pipeline (GCF_002204515.2_AaegL5.0_genomic.gff) were quantified using featureCounts (https://doi.org/10.1093/bioinformatics/btt656) with default parameters. Count data were transformed to TPM (transcripts per million) values using a custom Perl script. Details on libraries, alignment statistics, and gene expression estimates (expressed in transcripts per million; TPM) are provided as Supplementary Data 4–8.

### Identification of ‘collapsed’ and ‘merged’ gene models from AaegL3.5 to AaegL5.0

VectorBase annotation AaegL3.5 was compared to NCBI *Aedes aegypti* annotation release 101 on AaegL5.0 using custom code developed at NCBI as part of NCBI’s eukaryotic genome annotation pipeline. First, assembly-assembly alignments were generated for AaegL3 (GCA_000004015.3) x AaegL5.0 (GCF_002204515.2) as part of NCBI’s Remap coordinate remapping service, as described at http://www.ncbi.nlm.nih.gov/genome/tools/remap/docs/alignments. The alignments are publicly available in NCBI’s Genome Data Viewer (https://www.ncbi.nlm.nih.gov/genome/gdv/), the Remap interface, and by FTP in either ASN.1 or GFF3 format (ftp://ftp.ncbi.nlm.nih.gov/pub/remap/Aedes_aegypti/2.1/). Alignments are categorized as either ‘first pass’ (aka reciprocity=3) or ‘second pass’ (aka reciprocity=1 or 2). First pass alignments are reciprocal best alignments, and are used to identify regions on the two assemblies that can be considered equivalent. Second pass alignments are cases where two regions of one assembly have their best alignment to the same region on the other assembly. These are interpreted to represent regions where two paralogous regions in AaegL3 have been collapsed into a single region in AaegL5, or vice versa.

For comparing the two annotations, both annotations were converted to ASN.1 format and compared using an internal NCBI program that identifies regions of overlap between gene, mRNA, and CDS features projected through the assembly-assembly alignments. The comparison was done twice, first using only the first pass alignments, and again using only the second pass alignments corresponding to regions where duplication in the AaegL3 assembly has been collapsed. Gene features were compared, requiring at least some overlapping CDS in both the old and new annotation to avoid noise from overlapping genes and comparisons between coding vs. non-coding genes. AaegL5.0 genes that matched to two or more VectorBase AaegL3.5 genes were identified. Matches were further classified as collapsed paralogs if one or more of the matches was through the second pass alignments, or as improvements due to increased contiguity or annotation refinement if the matches were through first pass alignments (e.g. two AaegL3.5 genes represent the 5’ and 3’ ends of a single gene on AaegL5.0, such as SPR. Detailed lists of merged genes are in Supplementary Data 10–11.

### Comparison of alignment to AaegL3.4 and AaegL5.0

The AaegL5.0 transcriptome was extracted from coordinates provided in GCF_002204515.2_AaegL5.0_genomic.gtf. The AaegL3.4 transcriptome was downloaded from Vectorbase (https://www.vectorbase.org/download/aedes-aegypti-liverpooltranscriptsaaegl34fagz. Salmon (v0.8.2)^68^ indices were generated with default parameters, and all libraries described in Supplementary Data 4 were mapped to both AaegL3.4 and AaegL5 transcriptomes using ‘quant’ mode with default parameters. Mapping results are presented as Supplementary Data 9 and Fig. 1h.

### ATAC-Seq

The previously described ATAC-Seq protocol was adapted for *Ae. aegypti* brains^69^. Individual female brains were dissected in 1X PBS, immediately placed in 100µL ice-cold ATAC lysis buffer (10 mM Tris-Hcl, pH 7.4, 10 mM NaCl, 3 mM MgCl2, 0.1% IGEPAL CA-630), and homogenized in a 1.5 ml Eppendorf tube using 50 strokes of a Wheaton 1 ml PTFE tapered tissue grinder. Lysed brains were centrifuged at 400 g for 20 min at 4^o^C and the supernatant was discarded. Nuclei were resuspended in 52.5 µL1X Tagmentation buffer (provided in the Illumina Nextera DNA Library Prep Kit) and 5 µL was removed to count nuclei on a hemocytometer. 50,000 nuclei were used for each transposition reaction. The concentration of nuclei in Tagmentation buffer was adjusted to 50,000 nuclei in 47.5 µL Tagmentation buffer and 2.5 µL Tn5 enzyme was added (provided in the Illumina Nextera DNA Library Prep Kit). The remainder of the ATAC-Seq protocol was performed as described^69^. The final library was purified and size-selected using double-sided AMPure XP beads (0.6x, 0.7x). The library was checked on an Agilent Bioanalyzer 2100 and quantified using the Qubit dsDNA HS Assay Kit. Resulting libraries were sequenced 75 bp paired-end on an Illumina NextSeq500 platform at an average read depth of 30.5 million reads per sample. Raw fastq reads were checked for nucleotide distribution and read quality using FASTQC (http://www.bioinformatics.babraham.ac.uk/projects/fastqc) and mapped to the AaegL5 and AaegL3 versions of the *Ae. aegypti* genome using Bowtie 2.2.9(ref. ^70^). Aligned reads were processed using Samtools 1.3.1(ref. ^71^) and Picard 2.6.0 (http://broadinstitute.github.io/picard/index.html), and only uniquely mapped and non-redundant reads were used for downstream analyses. To compare the annotation and assembly of the *sex peptide receptor* (*SPR*) gene in AaegL3 and AaegL5, we used NCBI BLAST^72^ to identify AAEL007405 and AAEL010313 as gene fragments in AaegL3.4 annotation that map to *SPR* in the AaegL5.0 genome (BLAST e-values for both queries mapping to *SPR* were 0.0). Next, we used GMAP^73^ to align AAEL007405 and AAEL010313 fasta sequences to AaegL5. The resulting .gff3 annotation file was utilized by Gviz^74^ to plot RNA-Seq reads and sashimi plot as well as ATAC-Seq reads in the region containing *SPR*. Transcription start site analysis was performed using HOMER 4.9 (ref. ^75^). Briefly, databases containing 2 kb windows flanking transcription start sites genome-wide were generated using the ‘parseGTF.pl’ HOMER script from AaegL3.4 and AaegL5.0 gff3 annotation files. Duplicate transcription start sites and transcription start sites that were within 20 bp from each other were merged using the ‘mergePeaks’ HOMER script. Coverage of ATAC-Seq fragments in transcription start site regions was calculated with the ‘annotatePeaks.pl’ script. Coverage histograms were plotted using ggplot2 2.2.1 in RStudio 1.1.383, R 3.4.2 (ref. ^53^).

### Chromosome fluorescent *in situ* hybridization

Slides of mitotic chromosomes were prepared from imaginal discs of 4^th^ instar larvae following published protocols^4,76,77^. BAC clones were obtained from the University of Liverpool^16^ or from a previously described BAC library^78^. BACs were plated on agar plates (Thermo Fisher) and a single bacterial colony was used to grow an overnight bacterial culture in LB broth plates (Thermo Fisher) at 37°C. DNA from the BACs was extracted using Sigma PhasePrep TM BAC DNA Kit (Sigma-Aldrich, catalogue #NA-0100). BAC DNA for hybridization was labelled by nick translation with Cy3-, Cy5-dUTP (Enzo Life Sciences) or Fluorescein 12-dUTP (Thermo Fisher). Chromosomes were counterstained with DAPI in Prolong Gold Antifade (Thermo Fisher). Slides were analysed using a Zeiss LSM 880 Laser Scanning Microscope at 1000x magnification.

### M locus analysis

#### Aligning chromosome assemblies and Bionano scaffolds

The boundaries of the M-locus were identified by comparing the current AaegL5 assembly and the AaegL4 assembly^5^ using a program called LAST^79^ (data not shown). To overcome the challenges of repetitive hits, both AaegL5 and AaegL4 assemblies were twice repeat-masked^63^ against a combined repeat library of TEfam annotated TEs (https://tefam.biochem.vt.edu/tefam/)^3^ and a repeatmodeler output^64^ from the *Anopheles* 16 genome project^80^. The masked sequences were then compared using BLASTN^72^ and we then set a filter for downstream analysis to include only alignment with ≥98% identity over 1000 bp. After the identification of the approximate boundaries of the M-locus (and m-locus), which contains two male-specific genes, *myo-sex*^15^ and *Nix*^14^, we zoomed in by performing the same analysis on regions of the M- and m-locus plus 2 Mb flanking regions without repeatmasking. In this and subsequent analyses, only alignment with ≥98% identity over 500 bp were included. Consequently, approximate coordinates of the M-locus and m-locus were obtained on chromosome 1 of the AaegL5 and AaegL4 assemblies, respectively. Super-scaffold_63 in the Bionano optical map assembly was identified by BLASTN^72^ that spans the entire M-locus and extends beyond its two borders.

#### Chromosome quotient (CQ) analysis

CQ^17^ was calculated for each 1000 bp window across all AaegL5 chromosomes. To calculate CQ, Illumina reads were generated from two paired sibling female and male sequencing libraries. To generate libraries for CQ analysis, we performed two separate crosses of a single LVP_AGWG male to 10 virgin females. Eggs from this cross were hatched, and virgin male and female adults collected within 12 hours of eclosion to verify their non-mated status. We generated genomic DNA from 5 males and 5 females from each of these crosses. Sheared genomic DNA was used to generate libraries for Illumina sequencing with the Illumina Tru-Seq Nano kit and sequencing performed on one lane of 150 bp paired-end sequencing on an Illumina NextSeq 500 in high-output mode.

For a given sequence S_i_ of a 1000 bp window, CQ_(Si)_=F_(Si)_/M_(Si)_, where F_(Si)_ is the number of female Illumina reads aligned to S_i_, and M_(Si)_ is the number of male Illumina reads aligned to S_i_. Normalization was not necessary for these datasets because the mean and median CQs of the autosomes (chromosomes 2 and 3) are all near 1. A CQ value lower than the 0.05 indicates that the sequences within the corresponding 1000 bp window had at least 20 fold more hits to the male Illumina data than to the female Illumina data. Not every 1000 bp window produces a CQ value because many were completely masked by RepeatMasker^63^. To ensure that each CQ value represents a meaningful data points obtained with sufficient alignments, only sequences with more than 20 male hits were included in the calculation.

The CQ values were then plotted against the chromosome location of the 1000 bp window (Fig. 2e). Under these conditions, there is not a single 1000 bp fragment on chromosomes 2 and 3 that showed CQ=0.05 or lower.

### 10X genomics library preparation and Illumina sequencing for analysis of structural variants (SV)

Two individual pupae, one male and one female, were selected from the first generation of inbreeding of the LVP_AGWG strain (Extended Data Fig. 1a). High molecular weight (HMW) genomic DNA was extracted using the Qiagen MagAttract kit according to manufacturer’s instructions with minor modifications (rapid vortexing was replaced by inversion and wide-bore pipette tips were used – both to prevent excessive shearing of DNA). DNA extracted from each individual pupae was loaded into a separate lane of a 10X Chromium instrument for barcode tagging of the amplicons, then an Illumina sequencing library was prepared. Each library was sequenced in duplicate on two lanes of an Illumina HiSeq 2500. Due to the potential for transposable elements to give false positive SV calls, the AaegL5 genome was hard masked using Repeatmasker 4.05 (ref. ^63^) using the Aaeg-Liverpool V1 repeat library. Unplaced primary scaffolds and secondary haplotypes (i.e. any scaffolds or contigs except chromosomes 1, 2 and 3) were not used for alignment. Sequences were aligned to the reference using BWA via the LongRanger-Align function. Variants were called using GATK HaplotypeCaller (GATK version 3.5.0)^81^ and filtered for quality (QD > 5), strand bias (FS < 60) and read position (RankSum < 8). Only biallelic SNPs were used for phasing and subsequent analyses. The full Longranger-WGS pipeline was run on each sample (Longranger v.2.1.5) with memory overrides for both the SNP/INDEL phasing and SV calling stages required due to the high heterozygosity found in these samples. The pipeline was run with the pre-called VCF from the prior variant calling ensuring that the same sites were genotyped and phased in all samples. A second SV calling pipeline, GROCSVs, was run on the BWA alignments generated for variant calling. Repeat regions detected by RepeatMasker were blacklisted ensuring that no SVs would be called within these regions. SVs were compared between each pair of technical replicates and both methods. SVs were compared based on position and merged if they showed a 95% pairwise overlap. Only structural variants that were found in both technical replicates for a sample were reported (Fig. 2f and Supplementary Data 13).

### Identification of *Ae. aegypti Hox* genes and *Hox* sequence alignment

*Ae. aegypti Hox* cluster (HOXC) genes were identified by utilizing BLASTP2.6.1+ (ref. ^82^) to search the *Ae. aegypti* genome for genes with high similarity to *D. melanogaster* HOXC genes. The identity of *Aedes* HOXC genes was further resolved by comparing the relative position of candidate genes within the HOXC. The sequences of HOXC genes in *D. melanogaster* (annotation version R6.17) and *D. virilis* (annotation version R.106) were retrieved from Flybase, www.flybase.org^83^. The sequences of HOXC genes in *An. gambiae* (PEST annotation, version AgamP4.4) were retrieved from VectorBase, www.vectorbase.org^84^. Predicted coding exons for all *Hox* genes were aligned with the full HOXC genomic region using GenePalette www.genepalette.org^85^, then each species HOXC were proportionally adjusted to scale in Adobe Illustrator. The tandem repeats adjacent to *pb* were identified using GenePalette to search for regularly-spaced sequences. Six identical 749 bp tandem repeats were discovered on the end of Chromosome 1R in *Ae. aegypti*, related to telomere-associated sequences in species without telomeres^28^. Similar repeats of 556 bp were found at the same position at the tip of chromosome 1q in *Cx. quinquefasciatus* genome assembly CpipJ3 (ref. ^5^). To compare the Hox-Exd interaction motifs, Hox protein sequences were aligned using Clustal-Omega^86^ (Fig. 2g, Extended Data Fig. 3a, and Supplementary Data 14).

### Identification and analysis of *Ae. aegypti* GST and P450 genes and validation of the repeat structure of the GSTe cluster

Genes were initially extracted from the AaegL5.0 genome annotation (NCBI Release 101) by text search and filtered to remove “off target” matches (e.g. “cytochrome P450 reductase”), then predicted protein sequences of a small number of representative transcripts used to search the protein set using BLASTp, to identify by sequence similarity sequences not captured by the text search (resulting in two additional P450s, no GSTs). For each gene family, predicted protein sequences were used to search the proteins of the AaegL3.4 gene set using BLASTp. All best matches, and additional matches with amino acid identity >90% were tabulated for each gene family (Supplementary Data 15) to identify both closely related paralogues and alleles annotated as paralogues in AaegL3.4. Based on BLASTp search against the AaegL3.4 protein set, the two putative P450 genes not annotated as such in AaegL5.0 (encoding proteins XP_001649103.2 and XP_021694388.1) appear to be incorrect gene models in the AaegL5.0 annotation, which should in fact be two adjacent genes (CYP9J20 and CYP9J21 for XP_001649103.2; CYP6P12 and CYP6BZ1 for XP_021694388.1). Compared to AaegL3.4, which predicts a single copy each of *GSTe2*, *GSTe5*, and *GSTe7*, the NCBI annotation of AaegL5.0 predicts three copies each of *GSTe2* and *GSTe5*, and 4 copies of *GSTe7*, arranged in a repeat structure. BLASTn searches revealed one additional copy each of *GSTe2* and *GSTe5* in the third duplicated unit. Both contain premature termination codons due to frameshifts, but these could be due to uncorrected errors in the assembly. Error correction of all duplicated units was not possible due to the inability to unequivocally align reads to units not ‘anchored’ to adjacent single-copy sequence. To validate these tandem duplications, one lane of whole genome sequence data from a single pupa of the LVP_AGWG strain (H2NJHADXY, lane 1) was aligned to the AaegL3 scaffold containing the GSTe cluster (“supercont1.291”) using bowtie2 v2.2.4 (ref. ^87^), with ‘--very-fast-local’ alignment parameters, an expected fragment size between 0 and 1500 bp and relative orientation “forward-reverse” (“-I 0 -X 1500 -fr”). Reads with a mapping quality less than 10 were removed using Samtools^71^. Read pairs aligned to either DNA strand (“-s 0”) and overlapping a gene’s exons by at least 100 bp (“--minOverlap 100”) were assigned to genes using ‘featureCounts’, part of the ‘Subread’ v1.5.0-p2 package^88^ and plotted in Fig. 3b. To visualize the sequence identity of the repeat structure in the GSTe cluster (Fig. 3c), we extracted the region spanning the cluster from AaegL5 chromosome 2 (351,597,324 - 351,719,186 bp) and performed alignment of PacBio reads using Gepard v1.4.0 (ref. ^89^). To validate this repeat structure, we aligned two *de novo* optical maps created by Bionano using linearized DNA labelled with Nt.BspQI or Nb.BssSI. Single molecules from both maps span the entire region and the predicted restriction pattern provides support for the repeat structure as presented in AaegL5 (Fig. 3d).

### Curation of proteases

First, we identified 404 genes annotated as serine proteases (proteases, proteinases, peptidases, trypsins and chymotrypsins) and metalloproteases (metalloproteases, metalloproteinases and metametallopetidases) in AaegL3.4, based on their conserved domains in AaegL3.4. The UniProt database was searched to confirm serine protease/metalloprotease molecular function. We mapped these transcripts against the AaegL5.0 transcriptome by taking the longest transcript and using the reciprocal best BLAST method. We then extracted the coding sequence (CDS) lengths and corresponding peptide lengths for each of the transcripts for each of the 404 genes, from both AaegL3.4 and AaegL5.0. This comparison showed that over 50% of the gene models corresponding to the two protease subclasses have been changed in AaegL5.0 (Supplementary Data 16). This does not include the change in the UTRs. Twenty-one of the previous models have been discontinued. We also analysed 49 more gene models that are annotated as serine proteases or metalloproteases in AaegL5.0 but not in AaegL3.4 and were able to map all of these back to AaegL3.4 gene models by reciprocal best BLAST. These genes were either not annotated or not identified as proteases in AaegL3.4.

### Curation of opsins and biogenic amine binding G protein-coupled receptors

Genes for the opsin and biogenic amine binding Class A G protein-coupled receptor (GPCR) superfamily were identified by tBLASTn searches against the *Ae. aegypti* AaegL5 genome assembly and manually annotated as previously described using multiple online databases and software^3,36,84,86,90–94^. The resulting gene models were assigned to putative functional classes on the basis of sequence homology to multiple vertebrate and invertebrate GPCRs that have been functionally characterized. Results are summarized in Extended Data Fig. 3b and Supplementary Data 17–19. In all, genes for 10 opsin and 17 biogenic amine-binding receptors were annotated [3 dopamine; 8 serotonin: 2 muscarinic acetylcholine; 3 octopamine/tyramine receptors; 1 “unclassified” Class A biogenic amine binding]. The AaegL5 assembly enabled the first full-length gene model predictions for two opsin (*GPRop10* and *12*) and 14 biogenic amine binding (*GPRdop1* and *2*, *GPR5HT1A*, *1B* and *2*, *putative 5HT receptor 1–3*, *GPRmac1* and *2*, *GPRoar 1, 2* and *4*, and *GPRnna19*) GPCRs, with consolidation of dopamine receptors from six to three, serotonin receptors from 11 to eight, muscarinic acetylcholine receptors from three to two, and octopamine/tyramine receptors from six to three by fusion and resolution of the AaegL3-derived models described in Nene et al., 2007^3^.

### Chemosensory receptors

#### Annotation of previously identified genes

We used previously published *Ae. aegypti* (hereafter *Aaeg*) odorant receptor (*OR*), gustatory receptor (*GR*), and ionotropic receptor (*IR*) sequences^95–97^as queries to locate these genes in the new assembly. TBLASTN^98^ was used for protein sequences and discontinuous MegaBLAST^99^ followed by GMAP^73^ was used for coding sequences. New gene models were built, or NCBI RefSeq models accepted/modified, in the corresponding locations in a WebApollo v2 browser^94^.

Most modifications of RefSeq or previously published models were based on supporting RNA-Seq data^38^. These RNA-Seq data were derived from key chemosensory tissues in adult males and females and were loaded into WebApollo as short read alignments for each tissue (raw data available in the NCBI SRA database) and as alignments of *de novo* assembled transcripts prefiltered for those with TBLASTN homology to published chemoreceptors (TBLASTN e-value <10^−10^; *de novo* transcriptome available in the NCBI TSA database). We paid particular attention to pre-existing genes that were recognized by previous authors as fragments or were simply outside the normal length range for receptors in each of the three families. In most cases, we were able to extend these genes to full-length using GENEWISE^100^ with closely related receptor proteins as queries or by manual assessment of TBLASTN homology of flanking sequences to related receptors. For *ORs* and *IRs*, we used a reciprocal discontinuous MegaBLAST^99^ to verify that the coding sequences we annotated in AaegL5 corresponded to specific previously identified genes. This was necessary due to varying levels of sequence divergence between alleles found in AaegL3 and AaegL5. Moreover, we found many cases where two previously identified genes from AaegL3 mapped to the same locus in AaegL5, likely representing alternative haplotypes erroneously included on separate contigs in AaegL3 (classified as ‘merged’ genes in Fig. 4b and Supplementary Data 20). We note that although no geneset annotation exists, AaegL4^5^, which de-duplicated and scaffolded AaegL3 onto chromosomes, could be used to confirm these ‘merges’ as well.

#### Search for new genes in AaegL5

We also searched AaegL5 for new genes using the same TBLASTN results used to locate previously known genes (searches of AaegL5 with known chemoreceptor proteins). For new *ORs*, we manually examined all TBLASTN hits with an e-value cutoff below 10^−10^ after filtering for overlap with previously annotated genes using BedTOOLS^101^. For new *IRs*, we did the same but lowered the e-value cutoff to 10^−50^ to exclude ionotropic glutamate receptors (*iGluRs*), which have significant homology to *IRs*^97^. This approach identified a handful of new *IR* genes that were then used to query the *An. gambiae* genome (AgamP4) with a much more liberal TBLASTN e-value threshold of 1000 (*iGluRs* were ignored). Resulting discoveries in *An. gambiae* were then used to requery AaegL5 with the same liberal threshold and so forth in an iterative process until no new hits were identified. For *GRs*, we used proteins from *An. gambiae* ^102,103^ and *D. melanogaster*^104,105^ as TBLASTN queries in addition to pre-existing *Ae. aegypti* proteins, and manually examined any hits with e-values below 1000. For all three families, instances of apparent loss of an *Ae. aegypti* chemoreceptor suggested by the tree analyses were checked by searching the NCBI transcriptome shotgun assembly database for *Ae. aegypti* with TBLASTN using the relevant *Cx. quinquefasciatus*, *An. gambiae*, or *D. melanogaster* protein as query. For many chemoreceptors, these searches of the transcripts should allow detection of more divergent proteins because they are longer than the shorter exons in the genome and independent of the genome assembly. However, no new genes were discovered by searches of the transcripts that had not been found in the genome, indicating that our compilations of these three chemoreceptor families are likely exhaustive. We checked whether newly identified genes were missing in the old assembly or present but simply not recognized as receptors/genes. We used BLASTN^98^ to query the AaegL3 assembly with the new receptor genes and BedTOOLS^101^ to exclude hits to previously identified receptors. We considered a gene to be present, fragmented, or missing if this approach revealed full-length homologous sequences (every exon in order on same contig), partial homologous sequences (only some exons or exons on two different contigs), or no homologous sequences, respectively.

#### Search for new genes in *An. gambiae* genome

Our search for new *IRs* in AaegL5 involved an iterative process that resulted in the discovery of ~60 new *IR* genes in *An. gambiae*. We also used new *GR* genes in AaegL5 to uncover 4 new *GRs* in *An. gambiae*. These models were built in the WebApollo instance at VectorBase and will be available in future updates to the *An. gambiae* gene set. Putative phylogenetic relationships and protein sequences for these new *An. gambiae* genes can be found in Fig. 4c, Extended Data Fig. 4, and Supplementary Data 23).

#### ‘Corrected’ and ‘fixed’ genes

A significant minority of genes in the AaegL5 assembly contained loss-of-function (LOF) mutations that we inferred to be the result of either sequence/assembly errors or segregating polymorphism within the genome reference strain. In these cases, we chose to incorporate the intact alleles into our annotation set and analyses and refer to the genes as ‘corrected’ (minor updates to in-frame stop codons or small indels) or ‘fixed’ (major updates such as removal of large insertions of repetitive DNA or addition of a missing N-terminus). Sequence/assembly errors were inferred when both (1) LOF mutations occurred in regions where alignments of short-read Illumina data from the reference strain to AaegL5 were unusually spotty or showed sudden drops in coverage, and (2) we were able to find intact transcripts in a *de novo* transcriptome^38^. The short-read Illumina data was included as a supporting track in the WebApollo instance used for annotation. Simple polymorphic LOF mutations such as in-frame stop codons and small frameshifting indels were also obvious in the aligned short-read Illumina data. Large polymorphic insertions of repetitive DNA were harder to detect, but were also ‘fixed’ when we were able to find intact alleles in either the NCBI TSA database or the previous AaegL3 genome assembly. Details on the types of LOF mutations corrected and the source of the intact sequences can be found in (Supplementary Data 20).

#### Chemosensory receptor gene naming

We chose to retain previously published names for all *ORs* and *GRs*, simply dropping one of the two pre-existing names for gene pairs that were merged into a single locus in AaegL5, and starting the numbering for new genes where the previous genesets left off. The only exceptions were a handful of *GR* isoforms that were renamed to maintain the standard of increasing lower case letter suffixes for sequentially ordered sets of private exons while accommodating new isoforms. The result for the *OR* and *GR* families is a set of non-sequential gene numbers with limited phylogenetic meaning but increased stability – a priority given the large number of previous publications on *OR* genes in particular. In contrast to the *ORs* and *GRs*, however, we chose to rename the majority of *IR* genes in *Ae. aegypti*. We made this decision because our annotation efforts doubled the size of the family for this mosquito and produced what we expect to be a nearly exhaustive compilation. We used the following set of rules for renaming *IR* genes. The first two rules maintain pre-existing names, while the last two result in significant changes. (1) We retained *D. melanogaster* names for highly conserved *IRs* with clear 1-to-1 orthology across insects. These include *AaegIr8a*, *AaegIr21a*, *AaegIr25a*, *AaegIr40a*, *AaegIr60a, AaegIr68a*, *AaegIr76b*, and *AaegIr93a*. (2) We retained *D. melanogaster* names for relatively conserved *IRs* with clear 1-to-2 orthology in *Ae. aegypti*, adding a ‘1’ or ‘2’ suffix for the two genes in *Ae. aegypti*. These include *Ir87a* (*AaegIr87a1* and *AaegIr87a2*) and *Ir31a* (*AaegIr31a1* and *AaegIr31a2*). (3) We retained *D. melanogaster* roots for *IR* clades that are clearly related to specific *D. melanogaster* genes but have undergone more extensive species-specific expansion. These include genes in the *DmelIr75a*-*d* clade, the *DmelIr7a-g* clade, the *DmelIr41a* clade, and the *Dmel100a* clade. The corresponding members of each clade in *Ae. aegypti* were given the *D. melanogaster* number root with a single lower-case letter suffix (i.e. *AaegIr75a-l*, *AaegIr7a-r*, *AaegIr41a-q*, and *AaegIr100a-d*). Note that the specific suffix given in *Ae. aegypti* does not imply orthology with the *D. melanogaster* gene of the same suffix. (4) We renamed all remaining genes in *Ae. aegypti* starting with *AaegIr101* and increasing by single integers up to *AaegIr172*. The vast majority of these genes (all but *AaegIr101*-*AaegIr104*) fall into massive species-specific expansions loosely related to taste receptors in the *DmelIr20a* clade^37^. Only 25 of these 72 genes had been previously identified and all had names in the range of Ir101 to Ir120). We similarly added many new *IR* genes to the previously described *An. gambiae* genesets^97,103^, and renamed the entire family in that species according to the same rules. Old names for all *Ae. aegypti* and *An. gambiae IRs* are in Supplementary Data 20 and in parentheses on the ID line of the supplementary fasta file (Supplementary Data 23).

#### Tree building

The *Ae. aegypti* chemoreceptors in each family were aligned with those from *An. gambiae*^80,103^ (incorporating updates to *Agam IR* and *GR* families from the current work) and *D. melanogaster*^104,105^ using ClustalX v2^106^. Chemoreceptors annotated from another Culicine mosquito with a publicly-available genome sequence, *Cx. quinquefasciatus*^22^, have multiple near identical sequences that in light of our experience with *Ae. aegypti* are almost certainly the result of misassembly of alternative haplotypes. We therefore chose not to include receptors from that species in the trees, though we note that although no geneset annotation currently exists, CpipJ3^5^ has de-duplicated and scaffolded the existing *Cx. quinquefasciatus* genome assembly onto chromosomes, and will likely be useful in resolving the chemoreceptor gene families in *Cx. quinquefasciatus*. For *ORs* and *GRs*, poorly aligned regions were trimmed using TrimAl v1.4^107^ with the “gappyout” option that removes most poorly aligned or represented sequences. The *IR* family contains proteins that vary substantially in the length and sequence of their N-terminal regions; so for this family the “strict” option was employed in TrimAl, which removed much of their N-terminal alignment. Maximum likelihood phylogenetic analysis was conducted using PhyML v3.0^108^ with default parameters. In Fig. 4c and Extended Data Fig. 4, support levels for nodes are indicated by the size of black circles, reflecting approximate Log Ratio Tests (aLRT values ranging from 0–1 from PhyML v3.0 run with default parameters). Trees were arranged and coloured with FigTree v1.4.2 (http://tree.bio.ed.ac.uk/software/figtree/).

#### Expression analyses

We reanalysed published RNA-Seq data^38^ to quantify chemoreceptor expression in neural tissues using the new geneset (for details of alignment see methods section entitled “Alignment of transcriptome data to AaegL5 and quantification of gene expression”). We converted our webApollo-generated GFF3 file into the GTF format and provided this GTF file as input to featureCounts^109^. We counted reads across coding regions only since RNA-Seq evidence for UTRs was inconsistent across genes and some UTRs appeared to contain repetitive sequences that introduced mapping artefacts into inferred expression levels. We excluded UTRs by specifying the CDS flag (-t CDS) for each gene (-g gene_id) with an intact open-reading frame. We did not annotate UTRs for pseudogenes and were therefore able to count reads across all exons (-t exon) for genes whose coding regions were disrupted by loss-of-function mutations. We pooled reads across replicate libraries derived from the same tissue and time point and used the previously computed neurotranscriptome-wide normalization factors to calculate TPM-normalized expression levels for each chemoreceptor in each tissue. For visualization, we log10-transformed TPM expression levels using a pseudocount of 1. Expression values are presented in Extended Data Fig. 5–8.

### QTL mapping of dengue virus vector competence

#### Mosquito crosses

A large F_2_ intercross was created from a single mating pair of field-collected F_0_ founders. Wild mosquito eggs were collected in Kamphaeng Phet Province, Thailand in February 2011 as previously described^43^. Briefly, F_0_ eggs were allowed to hatch in filtered tap water and the larvae were reared until the pupae emerged in individual vials. *Ae. aegypti* adults were identified by visual inspection and maintained in an insectary under controlled conditions (28±1°C, 75±5% relative humidity and 12:12 hr light-dark cycle) with access to 10% sucrose. The F_0_ male and female initiating the cross were chosen from different collection sites to avoid creating a parental pair with siblings from the same wild mother^110,111^. Their F_1_ offspring were allowed to mass-mate and collectively oviposit to produce the F_2_ progeny (Extended Data Fig. 10). A total of 197 females of the F_2_ progeny were used as a mapping population to generate a linkage map and detect QTLs underlying vector competence for dengue virus (DENV).

#### Vector competence

Four low-passage DENV isolates were used to orally challenge the F_2_ females as previously described^43^. Briefly, four random groups of females from the F_2_ progeny were experimentally exposed two virus isolates belonging to dengue serotype 1 (KDH0026A and KDH0030A) and two virus isolates belonging to dengue serotype 3 (KDH0010A and KDH0014A), respectively. All four virus isolates were derived from human serum specimens collected in 2010 from clinically ill dengue patients at the Kamphaeng Phet Provincial Hospital^43^. Because the viruses were isolated in the laboratory cell culture, informed consent of the patients was not necessary for the present study. Complete viral genome sequences were deposited into GenBank (accession numbers HG316481–HG316484). Phylogenetic analysis assigned the viruses to known viral lineages that were circulating in Southeast Asia in the previous years^43^. Each isolate was amplified twice in C6/36 (*Ae. albopictus*) cells prior to vector competence assays. Four- to seven-day-old F_2_ females were starved for 24 hr and offered an infectious blood-meal for 30 min. Viral titres in the blood meals ranged from 2.0 × 10^4^ to 2.5 × 10^5^ plaque-forming units per ml across all isolates. Fully engorged females were incubated under the conditions described above. Vector competence was scored 14 days after the infectious blood-meal according to two conventional phenotypes: (*i*) midgut infection and (*ii*) viral dissemination from the midgut. These binary phenotypes were scored based on the presence/absence of infectious particles in body and head homogenates, respectively. Infectious viruses were detected by plaque assay performed in LLC-MK2 (rhesus monkey kidney epithelial) cells as previously described^43^,^112^.

#### Genotyping

Mosquito genomic DNA was extracted using the NucleoSpin 96 Tissue Core Kit (Macherey-Nagel). For the F_0_ male, it was necessary to perform whole-genome amplification using the Repli-g Mini kit (Qiagen) to obtain a sufficient amount of DNA. F_0_ parents and females of the F_2_ progeny were genotyped using a modified version of the original double-digest restriction-site associated DNA (RAD) sequencing protocol^113^, as previously described^114^. The final libraries were spiked with 15% PhiX, and sequenced on an Illumina NextSeq 500 platform using a 150-cycle paired-end chemistry (Illumina). A previously developed bash script pipeline^114^ was used to process the raw sequence reads. High-quality reads (Phred scores >25) trimmed to the 140-bp length were aligned to the AaegL5 reference genome (July 2017) using Bowtie v0.12.7 (ref. ^70^). Parameters for the ungapped alignment included ≤3 mismatches in the seed, suppression of alignments with >1 best reported alignment under a “try-hard” option. Variant and genotype calling was done from a catalogue of RAD loci created with the ref_map.pl pipeline in Stacks v1.19 (ref. ^115,116^). Downstream analyses only used high-quality genotypes at informative markers that were homozygous for alternative alleles in the F_0_ parents (e.g., AA in the F_0_ male and BB in the F_0_ female), had a sequencing depth ≥10×, and were present in ≥60% of the mapping population.

#### Linkage map

A comprehensive linkage map based on recombination fractions among RAD markers in the F_2_ generation was constructed using the R package OneMap v2.0–3 (ref. ^117^). Every informative autosomal marker is expected to segregate in the F_2_ mapping population at a frequency of 25% for homozygous (AA and BB) genotypes and 50% for heterozygous (AB) genotypes. Autosomal markers that significantly deviated from these Mendelian segregation ratios (*p*<0.05) were filtered out using a χ^2^ test. Due to the presence of a dominant male-determining locus on chromosome 1, fully sex-linked markers on chromosome 1 are expected to segregate in F_2_ females with equal frequencies (50%) of heterozygous (AB) and F_0_ maternal (BB) genotypes, because the F_0_ paternal (AA) genotype only occurs in F_2_ males. As previously reported^18^, strong deviations from the expected Mendelian segregation ratios were observed for a large proportion of markers assigned to chromosome 1 in the female F_2_ progeny. Markers on chromosome 1 were included if they had heterozygous (AB) genotype frequencies inside the ]40% - 60%[ range and F_0_ maternal (BB) genotype frequencies inside the ]5% - 65%[ range. These arbitrary boundaries for marker selection were largely permissive for partially or fully sex-linked markers on chromosome 1. Due to a lack of linkage analysis methods that deal with sex-linked markers when only one sex is genotyped, the recombination fractions between all pairs of selected markers were estimated using the *rf.2pts* function with default parameters for all three chromosomes. The *rf.2pts* function that implements the expectation-maximization (EM) algorithm was used to estimate haplotype frequencies and recombination rates between markers^11^ under the assumption of autosomal Hardy-Weinberg proportions. Due to this analytical assumption, the estimates of centiMorgans (cM) distances could be over- or under-estimated for markers on chromosome 1. Markers linked with a logarithm-of-odds (LOD) score ≥11 were assigned to the same linkage group. Linkage groups were assigned to the three distinct *Ae. aegypti* chromosomes based on the physical coordinates of the AaegL5 assembly. Recombination fractions were converted into genetic distances in cM using the Kosambi mapping function^118^. Linkage maps were exported in the R/qtl environment^119^ where they were corrected for tight double crossing-overs with the *calc.errorlod* function based on a LOD cutoff threshold of 4. Duplicate markers with identical genotypes were removed with the *findDupMarkers* function. To remove markers located in highly repetitive sequences, RAD sequences were blasted against the AaegL5 assembly using BLASTn v2.6.0. Markers with >1 blast hit on chromosomes over their 140-bp length and 100% identity were excluded from linkage analysis.

#### QTL mapping

The newly developed linkage map was used to detect and locate QTL underlying the DENV vector competence indices described above. Midgut infection was analysed in all F_2_ females whereas viral dissemination was analysed only in midgut-infected females. The four different DENV isolates were included as a covariate to detect QTL x isolate interactions. Single QTL detection was performed in the R/qtl environment^119^ using the EM algorithm of the *scanone* function using a binary trait model. Genome-wide statistical significance was determined by empirical permutation tests, with 1,000 genotype-phenotype permutations of the entire data set.

### Curation of ligand-gated ion channels and larvacidal activity of agricultural and veterinary insecticides

Putative *Ae. aegypti* cys-loop ligand-gated ion channel subunits were initially identified by searching the 2007 *Ae. aegypti* AaegL3 genome with TBLASTn^72^ using protein sequences of every member of the *D. melanogaster* cys-loop ligand-gated ion channel superfamily. In many cases the subunit coding sequences were incomplete due to regions showing low levels of homology, in particular the N-terminal signal peptide sequence and the hyper-variable intracellular region between the third and fourth transmembrane domains. These subunit sequences were used to search the latest AaegL5 RefSeq data through BLAST analysis, which in many cases completed missing sequence information (Supplementary Data 24). The neighbour-joining method^120^ and bootstrap resampling, available with the Clustal X program^106^, was used to construct a phylogenetic tree, which was then viewed using TreeView^121^ (Extended Data Fig. 9a).

To measure the effect of insecticides on *Ae. aegypti* larval behaviour, 3–10 larvae were dispensed manually into each well of a 96-well plate. Insecticides (imidacloprid, triflumezopyrim (targeting nAChRs^122^), abamectin (targeting primarily GluCls^123,124^) and fipronil (targeting primarily GABARs^123,124^) were added to a range of concentrations from 10^−11^ to 10^−4^ M. Larvae were incubated in compounds for 4 hours. The plate was then transferred to a video monitoring system (Extended Data Fig. 9b) which consisted of an LED light source backlighting the 96-well plate and an Andor camera. Images of the whole plate were acquired using a MATLAB script. The normalised movement index is plotted against the concentration of each compound. The movement index was derived by calculating the variance of a movie of each well in a 96-well plate and counting the number of pixels whose variance exceeds the mean variance by more than 4 standard deviations. Motility was estimated using the following algorithm: 1) a pair of images was acquired, separated by 10 ms. 2) the first image is subtracted from the second image to obtain a difference image. 3) pixels in the difference image with a value less than zero are set to zero. 4) pixels whose value is greater than 3 standard deviations above the mean of the image are set to 1, the rest are set to 0. 5) The mean of the pixels in each well is calculated to give a movement index. 6) The movement indices for the entire plate is divided by the maximum value to yield a normalized movement index.

### Mapping insecticide resistance and VGSC

The mosquito population Viva Caucel from Yucatán State in Southern Mexico (Longitude -89.71827, Latitude 20.99827), was collected in 2011 by Universidad Autónoma de Yucatán. We identified up to 25 larval breeding sites from 3–4 city blocks and collected ~1000 larvae. Larvae were allowed to eclose, and twice a day we aspirated the adults from the cartons, discarding anything other than *Ae. aegypti*. 300–400 *Ae. aegypti* were released into a 2-foot cubic cage where they were allowed to mate for up to 5 days with *ad libitum* access to sucrose, after which they were blood fed to collect eggs for the next generation. 390 adult mosquitoes were then phenotyped for deltamethrin resistance. We exposed groups of 50 mosquitoes (3–4 days old) to 3 µg of deltamethrin-coated bottles for 1 hr. After this time, active mosquitoes were transferred to cardboard cups and placed into an incubator (28ᵒC and 70% humidity) for 4 hr; these mosquitoes were referred as the resistant group. Knockdown mosquitoes were transferred to a second cardboard cup. After 4 hr, newly recovered mosquitoes were aspirated, frozen, and labelled as recovered; these were excluded from the current study. The mosquitoes that were knocked down and remained inactive at 4 hr post-treatment were scored as susceptible. DNA was isolated from individual mosquitoes by the salt extraction method^125^ and resuspended in 150 µL of TE buffer (10 mM Tris-HCl, 1 mM EDTA pH 8.0). We constructed a total of four gDNA libraries. Two groups were pooled from DNA of 25 individual females that survived 1 hr of deltamethrin exposure (resistant replicates 1 and 2). The second set of two libraries was obtained by pooling DNA from 25 females that were knocked down and inactive at 4 hr post treatment (susceptible replicates 1 and 2). Before pooling, DNA from each individual mosquito was quantified using the Quant-IT Pico Green kit (Life Technologies, Thermo Fisher Scientific Inc.) and around ~40 ng from each individual DNA sample (25 individuals per library) was used for a final DNA pool of 1 µg. Pooled DNA was sheared and fragmented by sonication to obtain fragments between 300–500 bp (Covaris Ltd., Brighton, U.K.). We prepared one library for each of the four DNA pools following the Low Sample (LS) protocol from the Illumina TrueSeqDNA PCR-Free Sample preparation guide (Illumina, San Diego CA). Because 65% of the *Ae. aegypti* genome consists of repetitive DNA, we performed an exome-capture hybridization to enrich for coding sequences using custom SeqCap EZ Developer probes (NimbleGen, Roche). Probes covered protein coding sequences (not including UTRs) in the AaegL1.3 genebuild using previously specified exonic coordinates^126^. In total, 26.7 Mb of the genome (2%) was targeted for enrichment. TruSeq libraries were hybridized to the probes using the xGen^®^Lock^®^Down recommendations (Integrated DNA Technologies). The targeted DNA was eluted and amplified (10–15 cycles) before being sequenced on one flow cell of a 100 bp HiSeq Rapid-duo paired-end sequencing run (Illumina) performed by the Centers for Disease Control (Atlanta, GA, USA).

The raw sequence files (*.fastq) for each pair-ended gDNA library were aligned to a custom reference physical map generated from the assembly AaegL5. Nucleotide counts were loaded into a contingency table with 4 rows corresponding to Alive Rep1, Alive Rep2, Dead Rep1, and Dead Rep2. The numbers of columns (c) corresponded to the number of alternative nucleotides at a SNP locus. The maximum value for c is 6, corresponding to A, C, G, T, insert, or deletion. Three (2 × c) contingency tables were subjected to χ^2^ analyses (c-1 degrees of freedom) to determine if there are significant (p ≤ 0.05) differences between 1) Alive replicates, 2) Dead replicates, and 3) Alive vs. Dead. If analysis 1) or 2) was significant, then that SNP locus was discarded. Otherwise the third contingency table consisted of 2 rows corresponding to Alive (sum of Reps 1 and 2), Dead (Reps 1 and 2 summed), and c columns. The χ^2^ value from the (2 × c) contingency χ^2^ analysis with (c-1) degrees of freedom was loaded into Excel to calculate the one-tailed probability of the χ^2^ distribution probability (p). This value was transformed with –log_10_(p). The experiment–wise error rate was then calculated following the method of Benjamini and Hochberg^127^ to lower the number of Type I errors (false positives).

### Data availability statement

All raw data have been deposited at NCBI under the following BioProject Accession numbers: PRJNA318737 (Primary Pacific Biosciences data, Hi-C sequencing data, whole-genome sequencing data from a single male Fig. 3b, and pools of male and females Fig. 2b-e, Bionano optical mapping data Fig 2d and Fig 3d, and 10X Linked-Read sequences Fig. 2f and Supplementary Data 13); PRJNA236239 (RNA-seq reads and de novo transcriptome assembly^38^ Fig. 1h-i and Supplementary Data 4–5,7,9); PRJNA209388 (RNA-seq reads for developmental time points^67^ Fig. 1h and Supplementary Data 4–6,9); PRJNA419241 (RNA-Seq reads from adult reproductive tissues and developmental time points, Verily Life Sciences Fig. 1h and Supplementary Data 4–5,8–9); PRJNA393466 (full-length Pacific Biosciences Iso-Seq transcript sequencing); PRJNA418406 (ATAC-Seq data from adult female brains, Fig 1i-j); PRJNA419379 (whole-genome sequencing data from colonies Fig. 5a); PRJNA399617 (RAD-Seq data Fig. 5c-f); PRJNA393171 (exome sequencing data Fig. 5g-i). The genome assembly and annotation is available from NCBI’s Assembly resource under accession GCF_002204515.2.

---

**Extended Data Figure 1.**
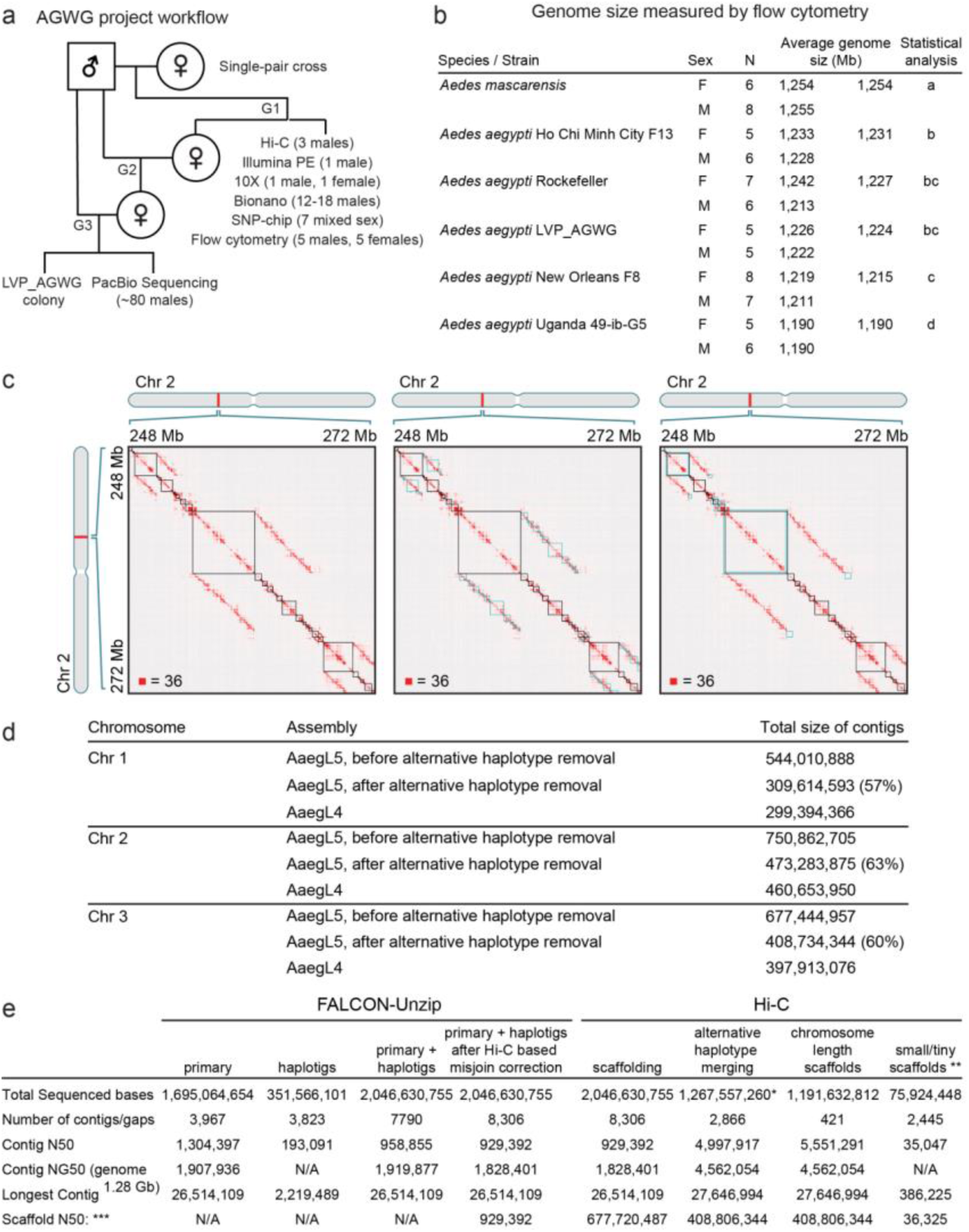
Project flowchart, measured genome size, and assembly process. **a**, Flowchart of LVP_AGWG strain inbreeding, data collection, and experimental design underlying the AaegL5 assembly process. **b**, Estimated average 1C genome size for each strain for 5 *Ae. aegypti* strains and *Ae. mascarensis*, the sister taxon of *Ae. aegypti* whose genome size has not previously been measured. There were no significant differences between the sexes within and between the species/strains analysed (p > 0.2). Significant differences between strains were determined using Proc GLM in SAS with both a Tukey and a Scheffé option with the same outcome. Data labelled with different letters are significantly different (p < 0.01). **c**, Combining Hi-C maps with 1D annotations enabled efficient review of sequences identified as alternative haplotypes by sequence alignment. Figure depicts a roughly 24 Mb x 24 Mb fragment of a contact map generated by aligning a Hi-C data set to a genome assembly generated during the process of creating AaegL5: a sequence comprising error-corrected, ordered and oriented FALCON-Unzip contigs. The intensity of each pixel in the contact map correlates with how often a pair of loci co-locate in the nucleus. Maximum intensity is indicated in the lower left of each panel. These maps include reads that do not align uniquely (reads with zero mapping quality). Because of the presence of highly similar sequences that represent alternative haplotypes, random assignment of reads to alternative haplotype loci by the aligner creates a characteristic off-diagonal signal that is easy to identify. Three panels show three types of annotations that are overlaid on top of the contact map: (left) Contig boundaries are highlighted as squares along the diagonal, (center) LASTZ-alignment-based annotations for contigs that are identified as alternative haplotypes to sequences incorporated into larger phased contigs. These contigs do not contribute sequence to merged chromosome-length scaffolds, (right) LASTZ-alignment-based annotations for contigs that only partially overlap in sequence with other contigs. These contigs contribute to the final fasta. **d**, Comparison of chromosome lengths between AaegL4 and AaegL5. Numbers are given prior to post-Hi-C polishing and gap closing. **e**, Step-wise assembly statistics for Hi-C scaffolding, alternative haplotype removal and annotation. *Removed length: 779,073,495 bp. See Dudchenko et al., 2017^5^ for definition of scaffold groups. Gaps between contigs were set to 500 bp for calculating scaffold statistics.

**Extended Data Figure 2.**
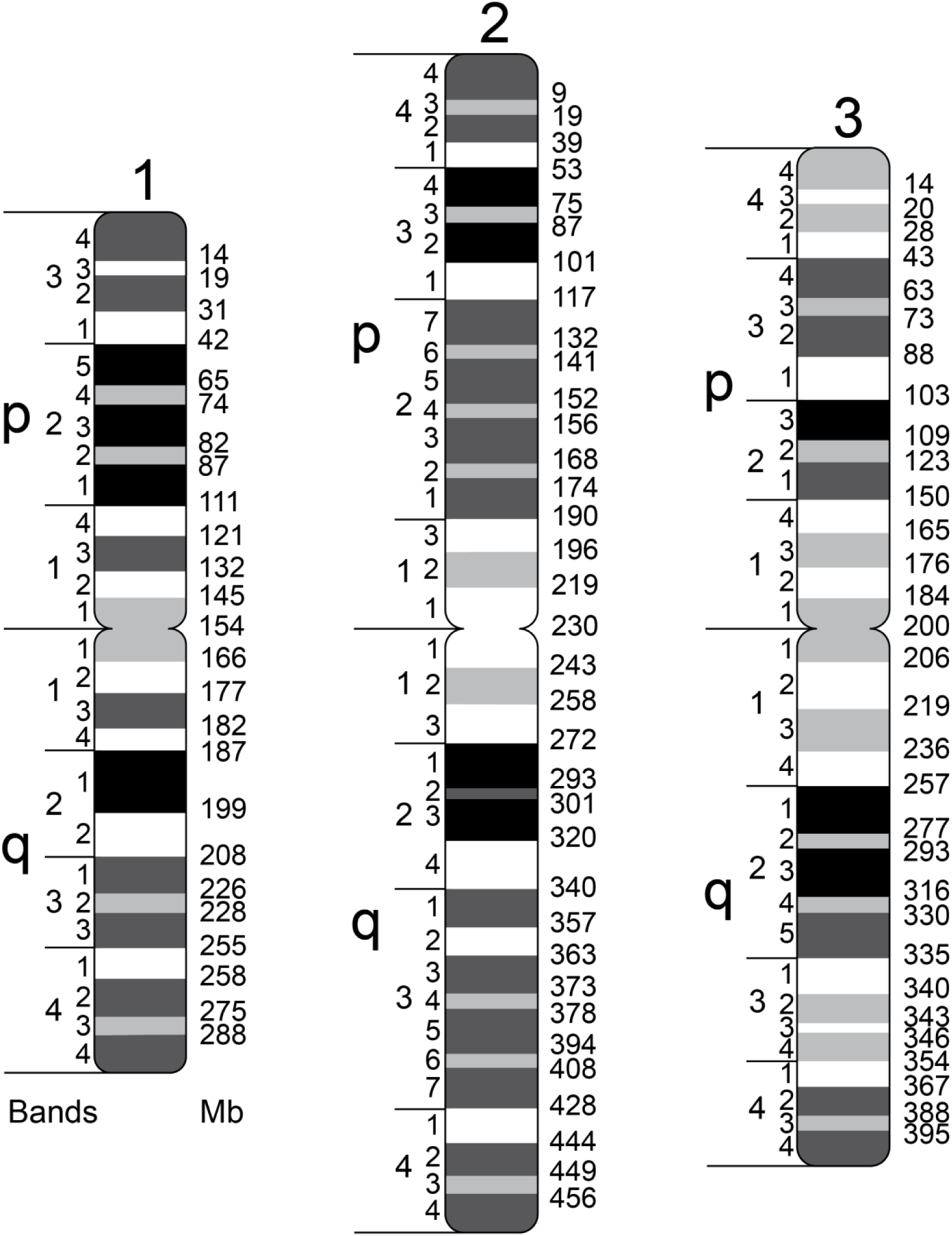
A chromosome map of the *Ae. aegypti* AaegL5 genome assembly. A physical genome map was developed by localizing 500 BAC clones to chromosomes using FISH. For the development of a final chromosome map for the AaegL5 assembly (Fig. 2a), we assigned the coordinates of each outmost BAC clone within a band (Supplementary Data 12) to the boundaries between bands. The final resolution of this map varies on average between 5 and 10 Mb because of the differences in BAC mapping density in different regions of chromosomes.

**Extended Data Figure 3.**
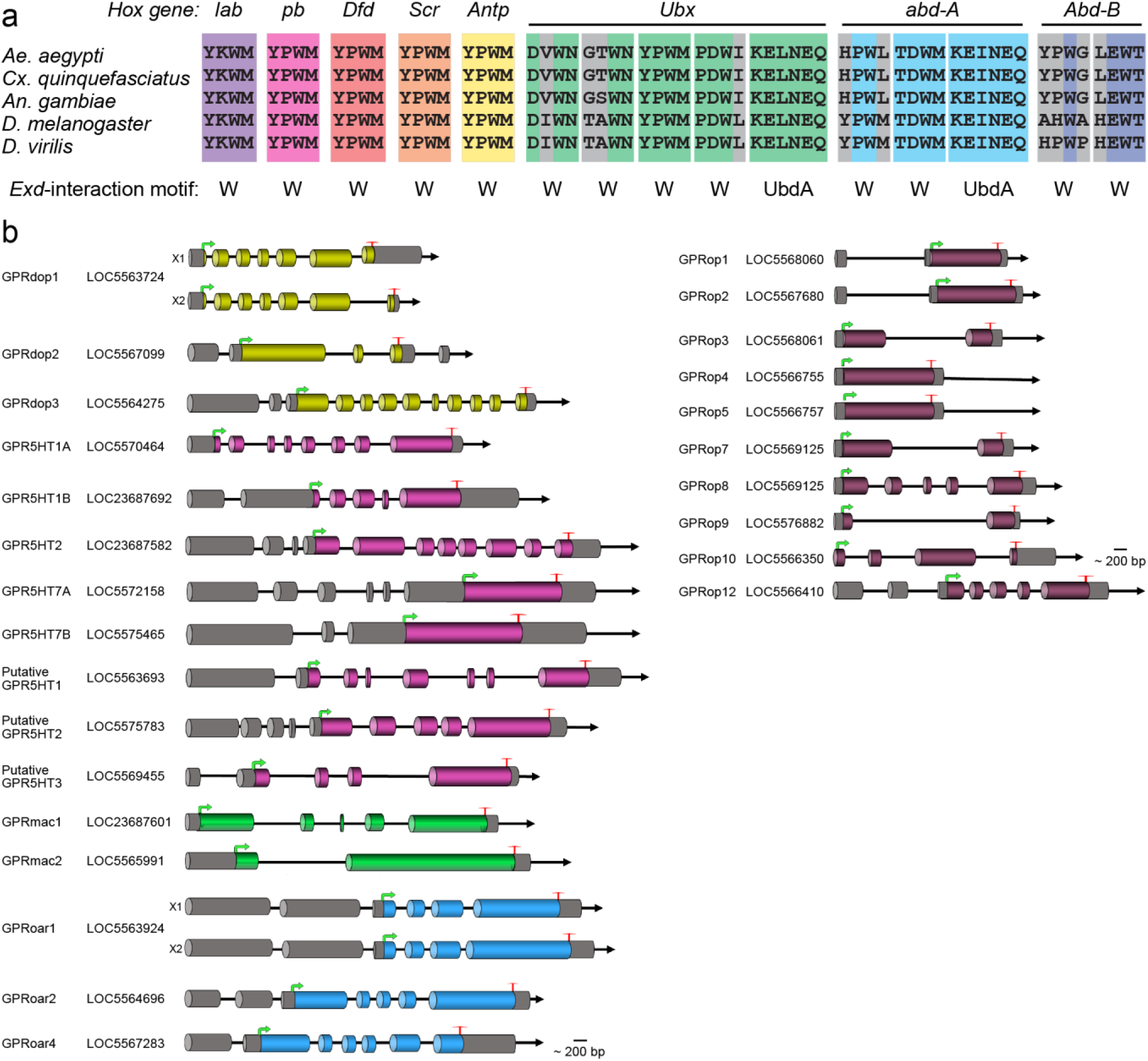
Hox interaction domains and GPCR gene models. **a**, Motifs known to mediate protein-protein interactions with the Hox cofactor Extradenticle (Exd)^128^ from the five indicated species are aligned using Clustal-Omega. Perfectly aligned residues are coloured according to Hox gene identity, non-conserved residues are grey. **b**, Schematic of predicted gene structures of the *Ae. aegypti* biogenic amine binding receptors (left) and opsins (right). Exons (cylindrical bars); introns (black lines); dopamine receptors (yellow bars); serotonin receptors (magenta bars); muscarinic acetylcholine receptors (green bars); octopamine receptors (blue bars); 5’ non-coding exons (dark shading). The “unclassified receptor” *GPRnna19* is not shown. Details on gene models compared to previous annotations and the predicted amino acid sequences of each gene are available in Supplementary Data 17–19.

**Extended Data Figure 4.**
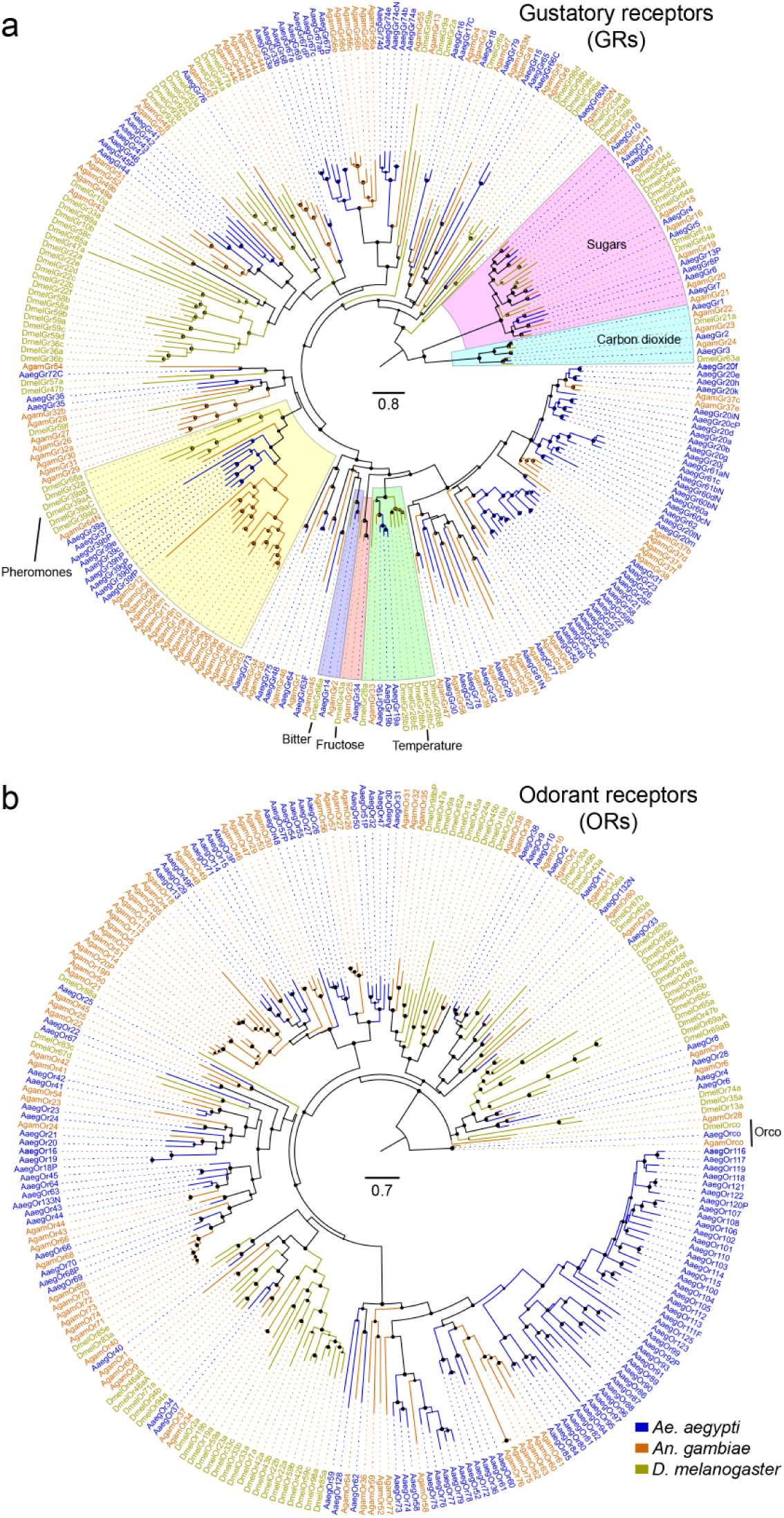
Phylogenetic trees of the gustatory receptor (*GR*) and odorant receptor (*OR*) gene families from *Ae. aegypti*, *An. gambiae*, and *D. melanogaster*. **a**, Maximum likelihood *GR* tree was rooted with the highly conserved and distantly related carbon dioxide and sugar receptor subfamilies, which together form a basal clade within the arthropod *GR* family^129^. Subfamilies and lineages closely related to *D. melanogaster GRs* of known function are highlighted. **b**, Maximum likelihood OR tree was rooted with Orco proteins, which are both highly conserved and basal within the *OR* family^104^. In panels **a-b**, support levels for nodes are indicated by the size of black circles – reflecting approximate Log Ratio Tests (aLRT values ranging from 0–1 from PhyML v3.0 run with default parameters^108^). Suffixes after protein names are C – minor assembly correction, F – major assembly modification, N – new model, and P – pseudogene. Scale bar: amino acid substitutions per site.

**Extended Data Figure 5.**
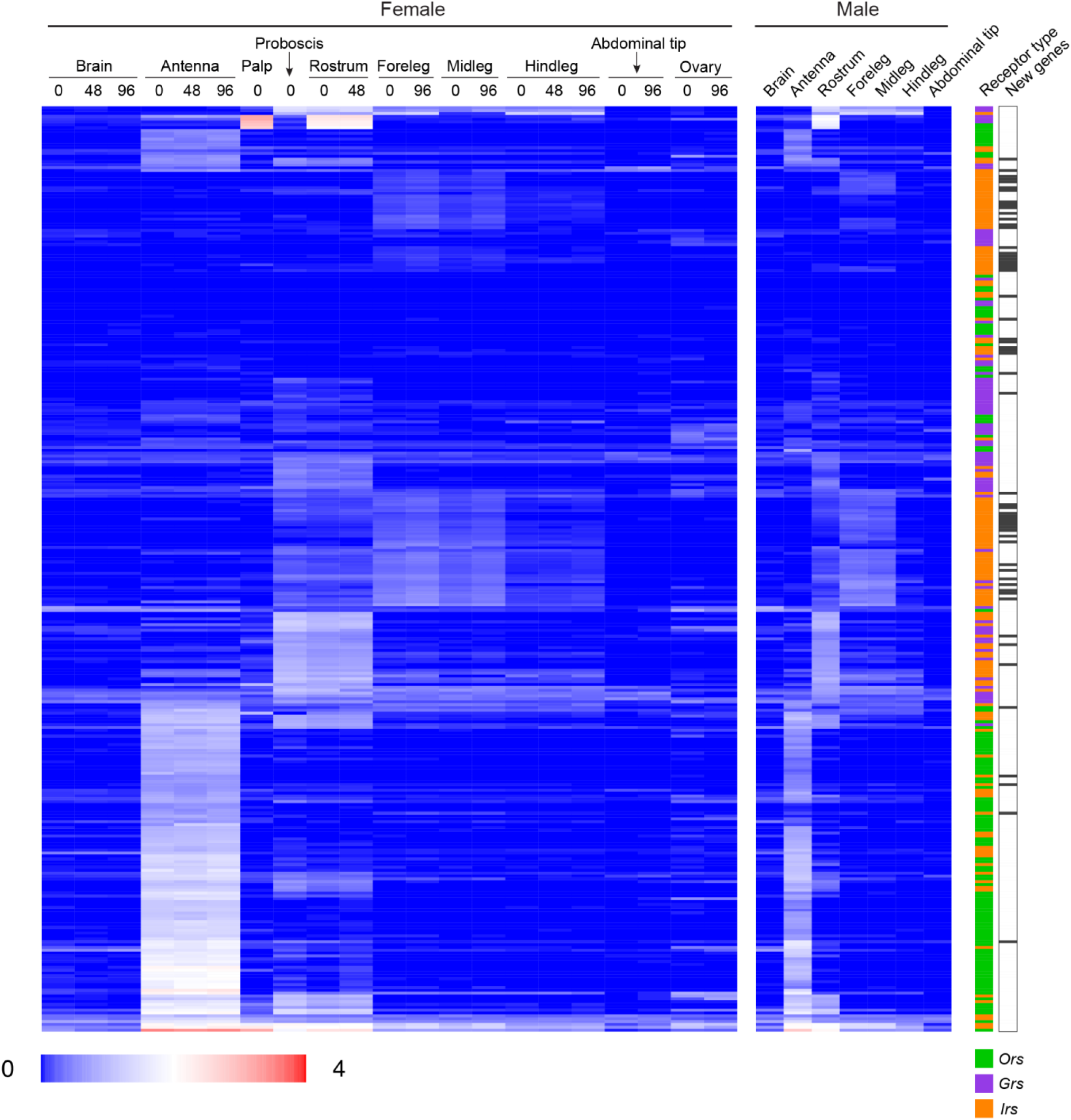
Chemosensory receptor expression in adult *Ae. aegypti* tissues. Previously published RNA-Seq data^38^ were reanalysed using the new chemoreceptor annotations and genome assembly. Expression is given for females in 3 stages of the gonotrophic cycle (0, 48, or 96 hr after taking a blood-meal, where 0 hr indicates not blood-fed, 48 hr indicates 48 hr after the blood-meal, and 96 hr indicates gravid). New genes are indicated by black bars at right.

**Extended Data Figure 6.**
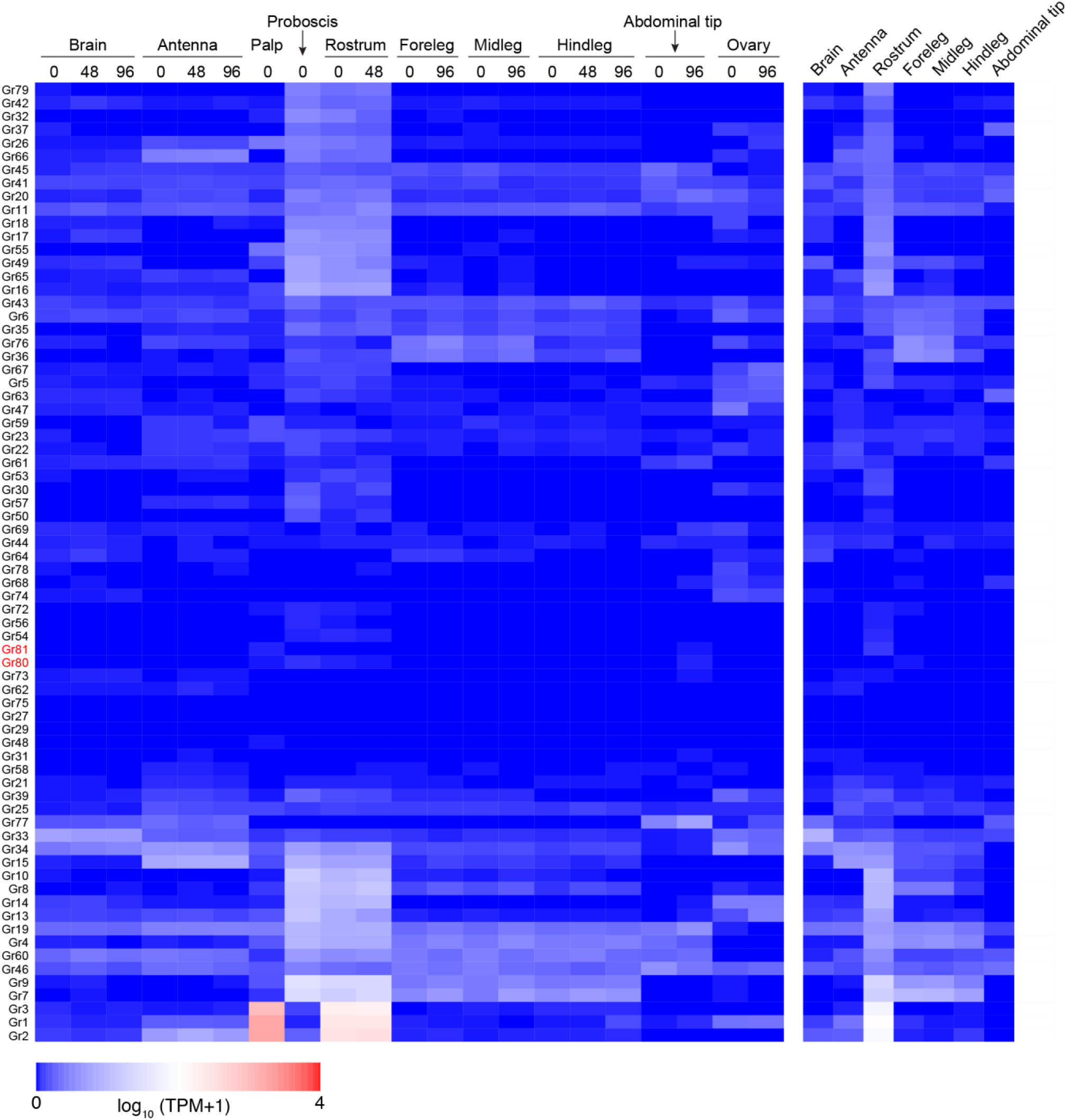
Gustatory receptor expression in adult *Ae. aegypti* tissues. Previously published RNA-Seq data^38^ were reanalysed using the new GR annotations and genome assembly. Expression is given for females in 3 stages of the gonotrophic cycle (0, 48, or 96 hr after taking a blood-meal, where 0 hr means not blood-fed). New genes are indicated by red text.

**Extended Data Figure 7.**
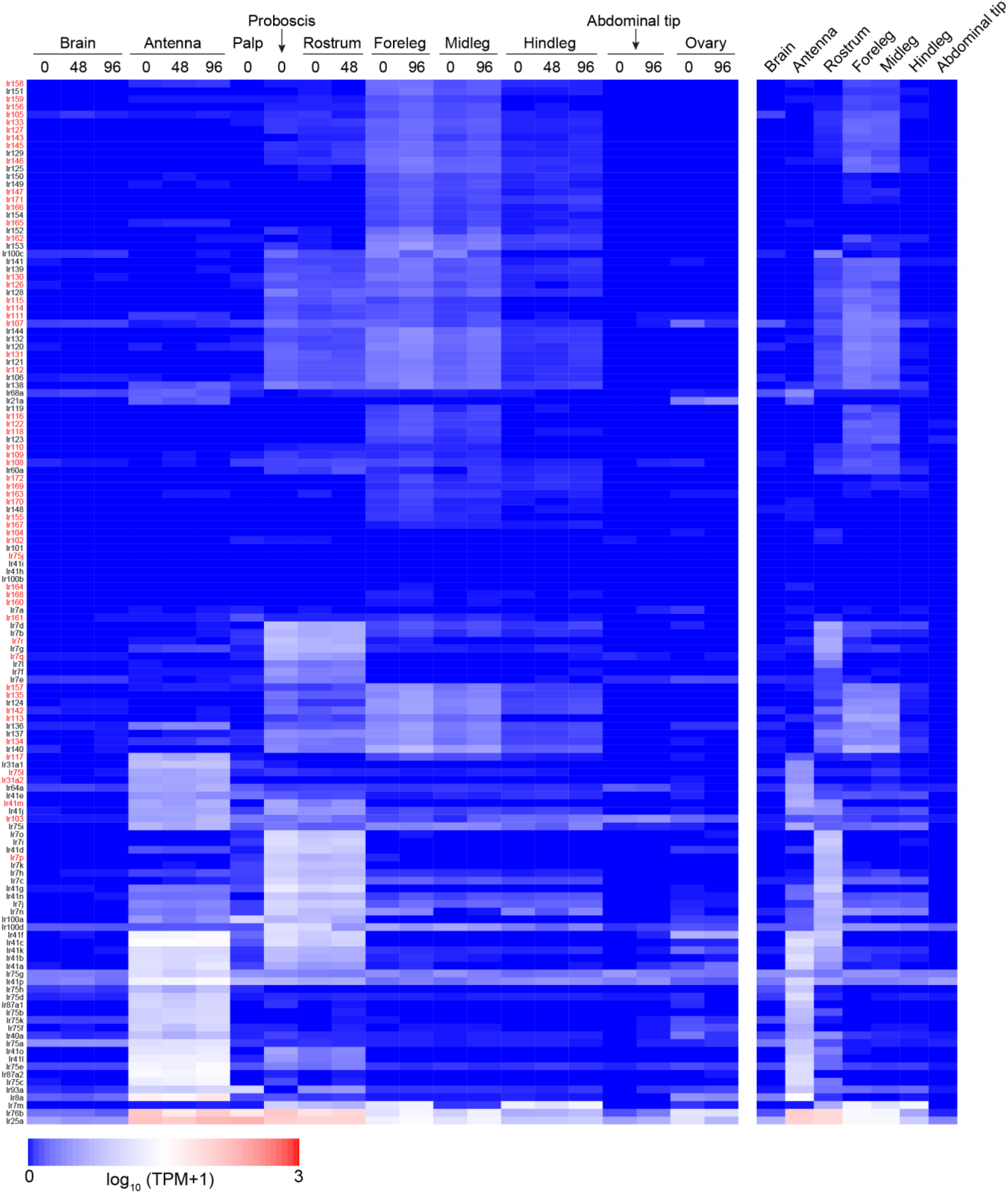
Ionotropic receptor expression in adult *Ae. aegypti* tissues. Previously published RNA-Seq data^38^ were reanalysed using the new IR annotations and genome assembly. Expression is given for females in 3 stages of the gonotrophic cycle (0, 48, or 96 hr after taking a blood-meal, where 0 hr means not blood-fed). New genes are indicated by red text.

**Extended Data Figure 8.**
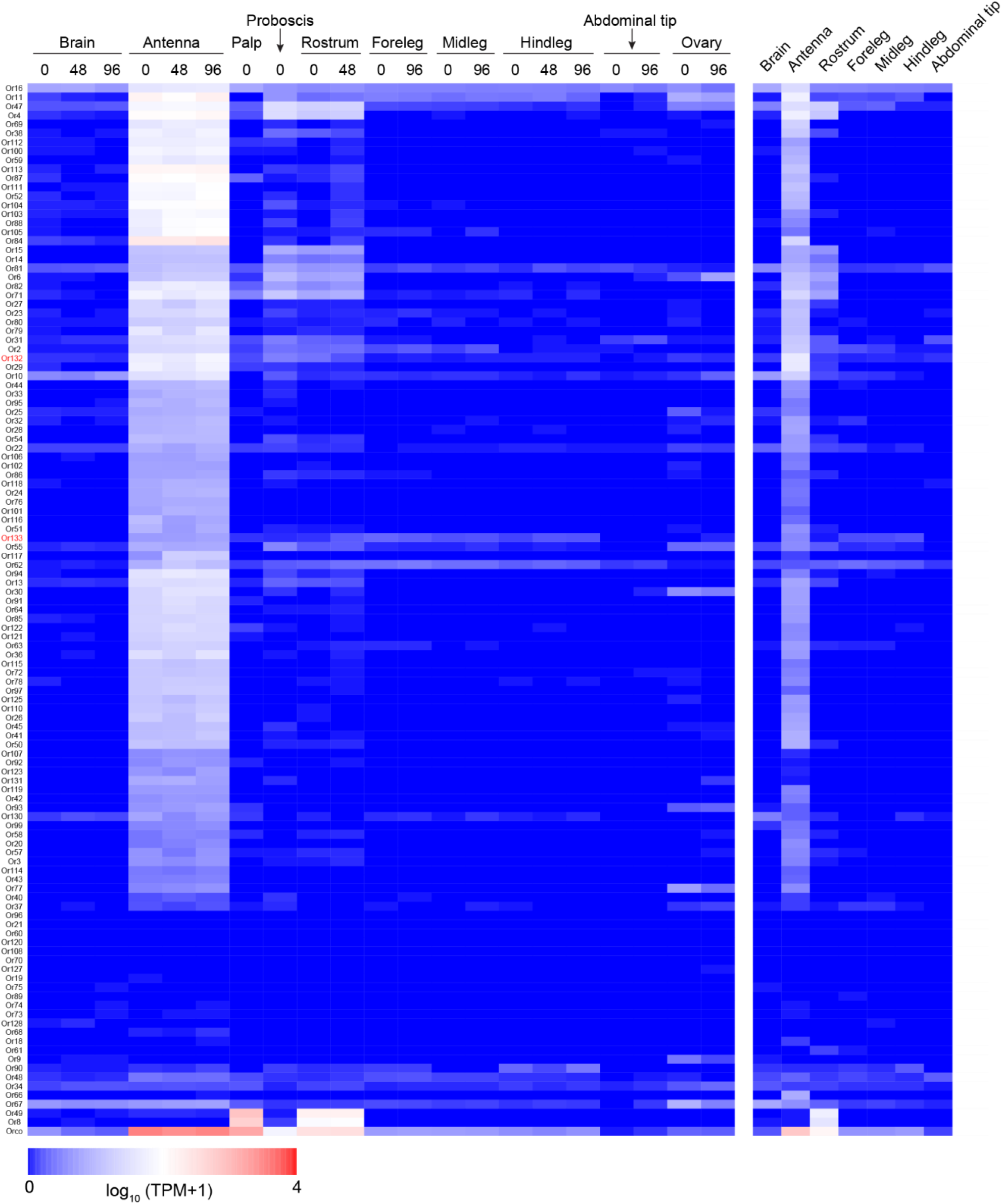
Odorant receptor expression in adult *Ae. aegypti* tissues. Previously published RNA-Seq data^38^ were reanalysed using the new OR annotations and genome assembly. Expression is given for females in 3 stages of the gonotrophic cycle (0, 48, or 96 hr after taking a blood-meal, where 0 hr means not blood-fed). New genes are indicated by red text.

**Extended Data Figure 9.**
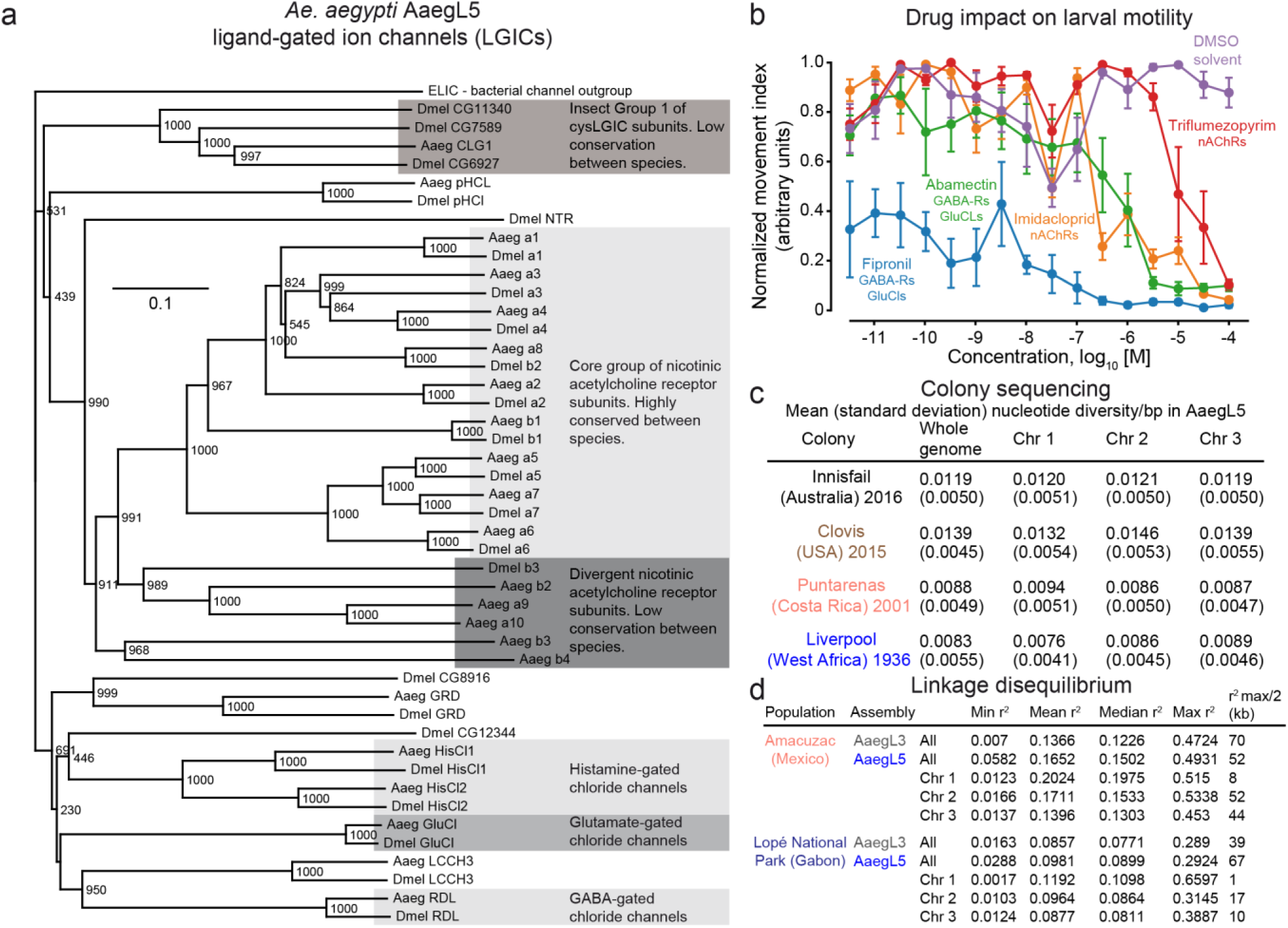
cys-loop ligand-gated ion channels and population genomic structure. **a**, Phylogenetic tree of cys-loop LGIC subunits for *Ae. aegypti* and *D. melanogaster*. The accession numbers of the *D. melanogaster* sequences used in constructing the tree are: Dα1 (CAA30172), Dα2 (CAA36517), Dα3 (CAA75688), Dα4(CAB77445), Dα5 (AAM13390), Dα6 (AAM13392), Dα7(AAK67257), Dß1 (CAA27641), Dß2 (CAA39211), Dß3 (CAC48166), GluCl (AAG40735), GRD (Q24352), HisCl1 (AAL74413), HisCl2 (AAL74414), LCCH3 (AAB27090), Ntr (NP_651958), pHCl (NP_001034025), RDL (AAA28556). For *Ae. aegypti* sequences see Supplementary Data 24. ELIC (Erwinia ligand-gated ion channel), which is an ancestral cys-loop LGIC found in bacteria (accession number P0C7B7), was used as an outgroup. Scale bar: amino acid substitutions per site. **b**, Concentration-response curves showing the impact on *Ae. aegypti* larval motility of insecticides currently used in veterinary and agricultural applications. **c**, Related to Fig. 5a. Mean nucleotide diversity in four colonized strains of *Ae. aegypti*, with standard deviation indicated in parentheses. Nucleotide diversity (π) was measured in non-overlapping 100 kb windows. **d**, Related to Fig. 5b. Table of linkage disequilibrium (r2) values along the *Ae. aegypti* AaegL5 genome assembly based on pairwise SNP comparisons. Data were obtained from the average r2 of SNPs in 1 kb bins.

**Extended Data Figure 10.**
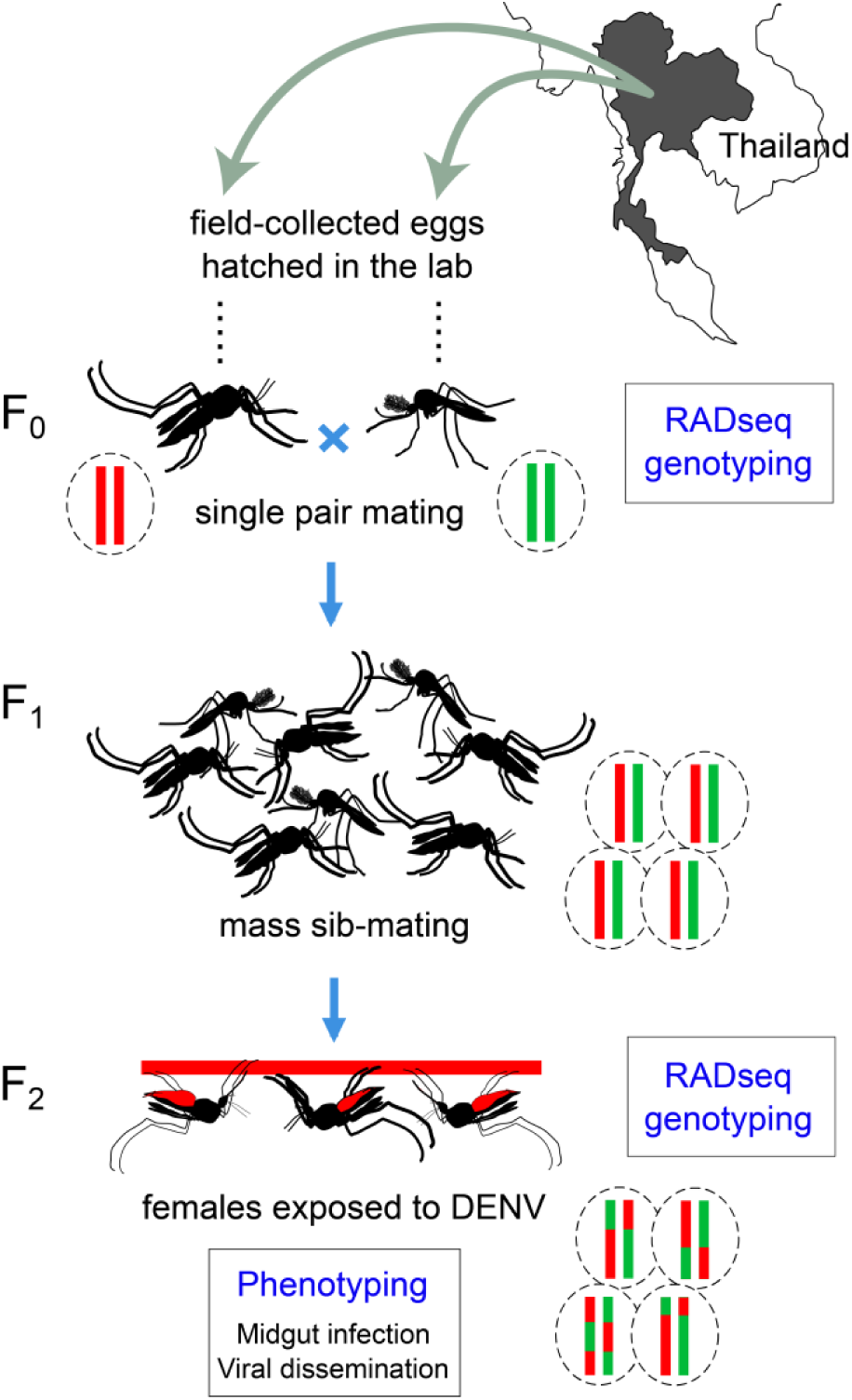
Schematic representation of the experimental workflow for Fig. 5d-f.

## Supplementary Information Guide

Files are available for download: http://www.dropbox.com/sh/93vlwt7wv9wspxa/AAAac1AxYP-8gca209KbdJyya?dl=0)

**Supplementary Data 1 - Fig 1e - TE - gff3 file with coordinates of TEs and repeats.gff**

Coordinates of identified TE and other repetitive sequences in GFF format

**Supplementary Data 2 - Fig 1e - TE - table of TE and repeat content.docx**

Data table listing the composition (in percent) of the AaegL5 genome assembly by class and family of transposable element and other repetitive sequences

**Supplementary Data 3 - Fig 1e - TE - repeat content by chromosome and scaffold.xlsx**

Data table listing the percentage of repetitive sequence content for each chromosome and unplaced scaffold

**Supplementary Data 4 - Fig 1f - gene expression - library information**

Library metadata for all RNA-Seq libraries aligned to AaegL5 genome

**Supplementary Data 5 - Fig 1f - gene expression - table of alignment statistics for all libraries.xlsx**

Alignment statistics for all RNA-Seq libraries aligned to AaegL5 genome

**Supplementary Data 6 - Fig 1f - gene expression - TPM Akbari.csv**

Gene expression in transcripts per million (TPM) for RNA-Seq libraries from developmental timepoints^67^. Reads were aligned to the AaegL5 genome and quantified using NCBI RefSeq Annotation version 101 (AaegL5.0)

**Supplementary Data 7 - Fig 1f - gene expression - TPM Matthews.csv**

Gene expression in transcripts per million (TPM) for RNA-Seq libraries from adult tissues^38^. Reads were aligned to the AaegL5 genome and quantified using NCBI RefSeq Annotation version 101 (AaegL5.0)

**Supplementary Data 8 - Fig 1f - gene expression - TPM Verily.csv**

Gene expression in transcripts per million (TPM) for previously unpublished RNA-Seq libraries (Verily Life Sciences) from developmental timepoints and adult reproductive tissues. Reads were aligned to the AaegL5 genome and quantified using NCBI RefSeq Annotation version 101 (AaegL5.0)

**Supplementary Data 9 - Fig 1h - L3 vs L5 table of alignment results.xlsx**

Table of pseudoalignment results (percent of reads mapped to annotated transcripts) for all RNA-Seq libraries for the AaegL3.4 and AaegL5.0 (RefSeq version 101) transcriptomes

**Supplementary Data 10- Fig 1 - merged paralogs.tsv**

NCBI GeneIDs and matched annotations from VectorBase AaegL3.5 for genes where multiple partially or fully redundant AaegL3.5 genes were collapsed into a single new gene annotation. Column 3 “is_best” designates the best matching VectorBase gene corresponding to each NCBI GeneID

**Supplementary Data 11 - Fig 1 - merged partial.tsv**

NCBI GeneIDs and matched annotations from VectorBase AaegL3.5 for genes where multiple non-overlapping AaegL3.5 genes were merged into a single new gene annotation. Column 3 “is_best” designates the best matching VectorBase gene corresponding to each NCBI GeneID

**Supplementary Data 12 - Fig 2a - BAC clone accessions and mapping locations.csv**

Positions of 88 BAC-ends mapped to the AaegL5 genome assembly and the position of the clones when hybridized to chromosomes

**Supplementary Data 13 - Fig 2f - Structural variants details.csv**

Summary table of structural variants identified by linked-read sequencing of two individual mosquitoes with analysis performed using Long Ranger and GROC-SVs software. Note that alignments were performed prior to standardizing gap length for the assembly at 100N; coordinates were subsequently transformed to the final NCBI reference by BLAST of surrounding regions

**Supplementary Data 14 - Fig 2g - AaegL5 HOXC gene table.xlsx**

Coordinates and summary of changes to genes in the HOX Cluster

**Supplementary Data 15 - Fig 3 - P450 and GST gene annotations.xlsx**

Coordinates and summary of changes to cytochrome P450 and GST gene families

**Supplementary Data 16 - Gene family annotations - Proteases.xlsx**

Coordinates and summary of changes to metalloproteases and serine genes

**Supplementary Data 17- Gene family annotations - Opsins and GPCRs Annotation Table.docx**

Coordinates and summary of changes to opsins and biogenic amine related genes

**Supplementary Data 18 - Gene family annotations - Biogenic amine peptide sequences.docx**

Peptide sequences of biogenic amine related genes. Non-synonymous substitutions between gene models predicted from the AaegL5 assembly compared to that predicted from the AaegL3 assembly (aqua shading); amino acid sequence unique to gene model predicted from the AaegL5 assembly (grey shading); amino acids associated with functional G protein-coupled receptors in other species (olive shading)

**Supplementary Data 19 - Gene family annotations - opsin peptide sequences.docx**

Peptide sequences of opsin genes. Non-synonymous substitutions between gene models predicted from the AaegL5 assembly compared to that predicted from the AaegL3 assembly (aqua shading); amino acid sequence unique to gene model predicted from the AaegL5 assembly (gray shading); amino acids associated with functional G protein-coupled receptors in other species (olive shading)

**Supplementary Data 20 - Fig 4 - Chemoreceptor Annotations Metadata 20171127.xlsx**

Metadata associated with each gene and alternatively spliced transcript in the AaegL5 geneset. Note that AaegL5 sequences for ‘fixed’ and ‘corrected’ genes have loss-of-function mutations that we inferred to be the result of within-strain polymorphism and/or sequence/assembly errors. We provide the coordinates for these genes in AaegL5 but used intact sequences (found in the NCBI TSA database, AaegL3, or apparent in short-read Illumina data from genome strain) in our analyses and fasta files. We also provide notes on the character and source of each LOF mutation for these genes

**Supplementary Data 21 - Fig 4 - Chemoreceptor Annotation.gff3**

Coordinates for gene models in gff3 format representing manual annotation of all chemoreceptors in the OR, IR, and GR gene families. Note that some genes are classified as pseudogenes here due to loss-of-function mutations that reflect sequencing errors or within strain polymorphism. The sequences for these genes were ‘fixed’ or ‘corrected’ in our analyses and in Supplementary Data 21 and 22

**Supplementary Data 22 - Fig 4 - ChemoreceptorCodingSequences.fasta**

Coding sequences for *Ae. aegypti* ORs, IRs, and GRs. Sequences modified relative to AaegL5 are indicated on the ID line as ‘corrected’ (minor assembly correction) or ‘fixed’ (major assembly correction). IRs and a few GR isoforms were renamed, and previously published names are included on the ID lines in parentheses

**Supplementary Data 23 - Fig 4 - ChemoreceptorPeptides.fasta**

Protein sequences for *Ae. aegypti* ORs, GRs, and IRs, and *An. gambiae* IRs (for which a significant number of new genes were identified). Sequences modified relative to AaegL5 are indicated on the ID line as ‘corrected’ (minor assembly correction), ‘fixed’ (major assembly correction), and ‘pseudogene’ (coding sequence repaired with stop codons coded as Z and other pseudogenizing mutations such as indels or splice site mutations coded as X). IRs and a few GR isoforms were renamed, and previously published names are included on the ID lines in parentheses

**Supplementary Data 24 - Gene family annotations - LGIC sequences and notes.docx**

Revised peptide sequences for *Ae. aegypti* cysLGICs subunits based on the AaegL5 sequence data. Lower case letters in the peptide sequences indicate the N-terminal signal peptide as predicted by SignalP 4.1

## SUPPLEMENTARY METHODS AND DISCUSSION

### FALCON configuration file

[General]

input_fofn = input.fofn

input_type = raw

length_cutoff = -1

genome_size = 1800000000

seed_coverage = 30

length_cutoff_pr = 1000

sge_option_da = -pe smp 5 -q bigmem

sge_option_la = -pe smp 20 -q bigmem

sge_option_pda = -pe smp 6 -q bigmem

sge_option_pla = -pe smp 16 -q bigmem

sge_option_fc = -pe smp 24 -q bigmem

sge_option_cns = -pe smp 12 -q bigmem

pa_concurrent_jobs = 96

cns_concurrent_jobs = 96

ovlp_concurrent_jobs = 96

pa_HPCdaligner_option = -v -B128 -t16 -M32 -e.70 -16400 -s100 -k18 -h480 -w8

ovlp_HPCdaligner_option = -v -B128 -M32 -h1024 -e.96 -12400 -s100 -k24

pa_DBsplit_option = -x500 -s400

ovlp_DBsplit_option = -s400

falcon_sense_option = --output_multi --min_idt 0.70 --min_cov 2 --max_n_read 200 --n_core 8

falcon_sense_skip_contained = True

overlap_filtering_setting = --max_diff 100 --max_cov 100 --min_cov 2 --n_core 12

### FALCON-Unzip assembly details

Half of the retained data were in reads of 16 kb or longer, with an average raw read length of 11.7 kb. We used raw reads 19 kb or longer as “seed reads” for error correction and generated 30.7 Gb (25.6X) of pre-assembled reads (preads) for contig assembly^6,56^. The resulting contig assembly contained primary contigs, comprising the backbone of the genome, and associated haplotigs, which represent phased, alternate haplotypes.

### Hi-C de-duplication

The Hi-C scaffolding procedure used both primary contigs and haplotigs from the FALCON-Unzip assembly as input, and here too undercollapsed heterozygosity was apparent. In fact, most genomic intervals were repeated, with variations, on 2 or more unmerged contigs, resulting in the ‘true’ genome length as measured by flow cytometry being far shorter than either the total length of the FALCON-Unzip contigs (2,047Mb), or the initial chromosome-length scaffolds (1,970Mb). The merge algorithm previously described^5^ detects and merges draft contigs that overlap one another due to undercollapsed heterozygosity. The parameters for running the merge have to be relatively permissive for AaegL5 to allow for identification of highly divergent overlaps separated by large genomic distances. We found that running the merge pipeline with such permissive parameters occasionally results in false positives and merges contigs that do not overlap. To avoid this, we developed a procedure to manually identify and ‘whitelist’ regions of the genome containing no overlap, based on both Hi-C maps and LASTZ alignments^130^.

The procedure consisted of surveying Hi-C maps with zero mapping quality reads while overlaying 2D annotations constructed from LASTZ output (see Extended Data Fig. 1c). This allows for two independent sources for confirming high sequence identity at two genomic intervals: one from short-read (mis)alignment and another one from contig sequence similarity. Based on the survey, the contig sequences representing chromosome-length scaffolds are split into “merge blocks” prior to creating the haploid reference: merge is allowed between contig sequences in the same blocks but not between contigs belonging to different blocks. This custom version of the merge pipeline is shared on GitHub (http://github.com/theaidenlab/AGWG-merge).

### Gap filling protocol

After Hi-C scaffolding and de-duplication, All 527 PacBio subread .fastq files were used as input to PBJelly for final polishing and gap-filling. The format for each input is denoted by the bold and italicized lines below (replace ***XXX_N.subreads.fastq*** with the full name of each file).

~~~
<jellyProtocol>
<reference>asm.fasta</reference>
<outputDir>./</outputDir>
<cluster>
<command notes=For SGE>echo ‘${CMD}’ | qsub -V -N “${JOBNAME}” -cwd -pe thread 8 -l mem_free=4G -o ${STDOUT} -e ${STDERR}</command>
<nJobs>100</nJobs>
</cluster>
<blasr>--minMatch 8 --minPctIdentity 70 --bestn 1 --nCandidates 20 --maxScore -500 --nproc 8 --noSplitSubreads</blasr>
<input baseDir=/seq/a_aegypti/pacbio/>
<job>XXX_1.subreads.fastq</job>
…
<job>XXX_527.subreads.fastq</job>
</input>
</jellyProtocol>
~~~

### BUSCO completeness

BUSCO, a benchmark based on single-copy universal orthologues^131^ was used to confirm that multiple haplotypes were present in the initial assembly and to evaluate the success of our de-duplication. BUSCO contains a database of genes which are thought to be present in single-copy in all species below a given taxonomic level. Thus, any complete assembly should include all or close to all of the genes. Since all these genes are also single-copy there should not be any duplicated genes in an assembly. Duplicates genes indicate potential alternate haplotypes present in the assembly result. BUSCO v1.22 was run with default parameters and the arthropoda gene set with the command:

BUSCO_v122.py –c 1 –f –in asm.fasta –o SAMPLE –l arthropoda –m genome

**Completeness of the final AaegL5 assembly** Complete: 2301 Single: 2133 Multi: 168 Fragment: 309 Missing: 65 Total BUSCO groups searched: 2675

**Completeness of the FALCON-Unzip assembly (primary contigs + haplotigs)** Complete: 2347 Single: 1101 Multi: 1246 Fragment: 274 Missing: 54 Total BUSCO groups searched: 2675

### Analysis of transposable elements

We ran RepeatMasker using the TEfam and Repbase databases, and found transposable elements represent 54.85% (excluding the 3.02% unclassified TEs) of the new assembled genome. Moreover, 25.48% of the total base pairs identified as TEs were DNA elements, 28.92% were RNA elements, and 0.45% were Penelope (Supplementary Data 1–3). Simple and tandem repeats occupy 3.3% of the genome, and the additional 7% consists of unclassified repetitive sequences. Similar to previous annotation, *Juan-A* in the Jockey family of non-long terminal repeat (non-LTR) retrotransposons accounts for ~3.4% of the genome, and is the most enriched TE type in the genome. Using Tandem Repeat Finder, we found that 6.9% of genome sequences are repeat sequences, while 1% of the genome is simple repeat sequences. Since a subset of the tandem repeat sequences overlap with TE regions, we then used tandem repeat finder to search for repeatmasked genome sequences and found that the whole genome contains 3.3% non-TE repetitive sequences.

We identified a significant positive correlation between GC content and the total lengths of TEs (Pearson’s r =0.37, p < 0.001) of each chromosome or scaffold. However, there is not a significant correlation between the number of TEs and GC content (Pearson’s r = -0.02, p > 0.05). Compared to previous TEfam annotation, new transposon elements such as CMC-Chapaev, CMC-Transib, sola, and Crypton showed relatively high copy numbers. Overall, a greater proportion of TE sequences belong to DNA elements compared to the previous annotation. Our results of TE identification using different libraries suggest that novel TE types are the main contributor to the higher proportion of DNA elements (Supplementary Data 1–3). However, it is difficult to directly compare these results with AaegL3 (ref. ^3^), since different TE elements may have different lengths and numbers of insertions, and many different element types have high sequence similarities.

### A split of the HOX-C gene cluster to two chromosomes

We examined the presence of motifs known to mediate protein-protein interactions with the Hox cofactor Extradenticle (Exd)^128^. Most Hox proteins bind Exd using the canonical “YPWM” motif, but in *D. melanogaster* the abdominal Hox proteins Ultrabithorax (Ubx) and Abdominal-A (Abd-A) have additional “W” motifs that may be utilized in a context-dependent manner^128^. The *Ae. aegypti* Hox proteins have all previously described “W” motifs (Extended Data Fig. 3a). In all three mosquito species analysed here, Abdominal-B (AbdB) has as an additional putative Exd interaction motif, “YPWG”, which closely resembles the canonical “YPWM” motif in other *D. melanogaster* Hox proteins (Extended Data Fig. 3a).

### Improved annotation of G protein-coupled receptors for vision and neuromodulation

Three dopamine receptors (*GPRdop1–3*) previously reported^3^ and subsequently characterized^93^, as well as eight putative serotonin (*GPR5HT*) receptors, one previously characterized^90^ (*GPR5HT7A*), two muscarinic acetylcholine receptors (*GPRmac1* and *2*) and three previously reported octopamine/tyramine receptors (*GPRoar1, GPRoar2*, and *GPRoar4*)^3,38^ were identified in the AaegL5 assembly and subsequently manually annotated. We made a considerable revision to *GPRdop3* with the addition of 241 amino acids to the 5’ region of the model and the inclusion of a short 11th exon (33 amino acids), increasing the total number of exons for this gene model from eight to 12.

The AaegL5 assembly enabled greater resolution of the serotonin or 5-hydroxytryptamine (5HT) receptor subfamily, comprising eight members (*GPR5HT1A, GPR5HT1B, GPR5HT2, GPR5HT7A, GPR5HT7B* and *putative 5HT receptors 1–3*) with prediction of the first full-length gene model for *GPR5HT1A* (80 amino acids added to the 5’ region) and substantial revision to *GPR5HT1B* (addition of 86 amino acids to the gene model, representing revision of sequence corresponding to exons 2, 5 and 6). The AaegL5.0 annotation also enabled major revision of *GPR5HT2* with the addition of 292 and 115 amino acids to the 5’ and internal regions of the model, respectively.

The remaining three receptors designated as *putative 5HT receptor 1–3* possess some sequence homology to vertebrate and invertebrate serotonin receptors. The AaegL3.4 gene models corresponding to these receptors comprise one or more exons with high amino acid similarity to vertebrate and invertebrate serotonin receptors, but lack 5’ and 3’ sequence and are considered incomplete. The revised AaegL5.0 gene models incorporated additional 5’ and/or 3’ sequence and each model comprises critical features inclusive of an initiation methionine, stop codon, seven transmembrane spanning domains, and canonical GPCR motifs such as N-terminal glycosylation and C-terminal palmitoylation sites. These three models are supported by RNA-Seq data and possess homology to orthologous serotonin receptors identified in vertebrate and invertebrate species. However, some ambiguity remains. The predicted protein sequence of the putative *GPR5HT receptor 2* is considerably longer than that of many GPCRs, for example, and the models will require resolution via molecular analyses.

Two muscarinic acetylcholine receptors (*GPRmac1* and 2) were identified using the AaegL5 assembly and possessed high amino acid identity to the AaegL3-derived models. The AaegL5 assembly also enabled greater resolution of the *Ae. aegypti* octopamine receptor subfamily (four complete gene models for *GPRoar1, GPRoar2*, and *GPRoar4*). Key advances include the prediction of two isoforms for *GPRoar1* (X1 and X2; addition of 149 and 141 amino acids to the 5’ region of X1 and X2, respectively) and the addition of a total 141 and 211 amino acids to the models for *GPRoar 2* and *4*, plus the deletion of 19 amino acids from *GPRoar4*.

Our analyses revealed several gene models for receptors that had been renamed between the AaegL3.4 and AaegL5.0 gene sets; specifically, *GPR5HT2* to a muscarinic M3 receptor (the original gene name was retained here) and *AaGPR5HT8* to an “uncharacterized protein” (subsequently renamed here as “*putative 5HT receptor 2*”). In the interest of consistency, the current analyses attempted to resolve these discrepancies (Supplementary Data 18) based on multiple lines of evidence, including RNA-Seq data, manual annotation, and sequence homology to functionally characterized GPCRs from vertebrates and invertebrates. Several instances of gene model collapse were identified between the AaegL3 and AaegL5 gene sets (AAEL014373 and AAEL017166 into LOC5564275 for *GPRdop3*; AAEL09573 and AAEL016993 into LOC5572158 for *GPR5HT7A*; AAEL015553 and AAEL002717 into LOC5575783 for *putative 5HT receptor 2*). We also note that multiple transcript variants were detected for *GPR5HT2* (LOC23687582; 6 variants), *GPR5HT7A* (LOC5572158; 4 variants) and *putative 5HT receptor 2* (LOC5575783; 14 variants). These variants were predicted to produce gene products with identical amino acid sequence and their status as haplotype sequence is yet to be resolved. Finally, one previously reported sequence (*GPRnna19*) that was identified as a putative biogenic amine binding receptor in the AaegL3.4 gene set was renamed in the AaegL5.0 gene set as a “putative tyramine/octopamine receptor”. The AaegL5.0 model includes an additional 142 amino acids of 5’ sequence, and is supported by RNA-Seq data and sequence homology to biogenic amine binding GPCRs. However, membrane prediction software suggests that this model comprise only three or four transmembrane spanning domains and the model lacks amino acid motifs considered critical to GPCR function. The gene was not assigned to subfamily in the present analysis. The model was designated as “unclassified” (Supplementary Data 18) and molecular studies will be needed to confirm the model. Finally, we note that the majority of GPCRs identified in the present analyses should be considered “orphan” receptors. Functional studies will be required for all but *GPRdop1, dop2*, and *5HT7A* to establish receptor activity and interaction with the cognate ligand.

Ten previously reported opsins (*GPRop1–5, GPRop7–10*, and *GPRop12*)^3,36^, were identified in the AaegL5 assembly, and sequence was confirmed via manual annotation. The opsins represent a gene family (typically 3–7 receptors in arthropods) of UV-, short- and long-wavelength sensitive receptors and have been annotated in many arthropods. Ten, 11 and 13 *opsin* genes were identified in the mosquitoes *Ae. aegypti, An. gambiae*, and *Cx. quinquefasciatus*, respectively^36^ and thus provide an opportunity to benchmark the AaegL5.0 annotation. All AaegL5.0 *opsins* were full-length and the predicted gene products possessed features indicative of functional GPCRs, including an initiation methionine, a stop codon, seven transmembrane domains, three extracellular and three intracellular loops, as well as conserved motifs associated with GPCR and opsin function (except for *GPRop10* which contained six transmembrane domains). Non-synonymous substitutions were identified in *AaegGPRop2, AaegGPRop7*, and *AaegGPRop10* in regions not typically considered critical for functions such as photon interaction, amine binding or G protein-coupling. AaegL5.0 enabled prediction of the 5’ coding region for *AaGPRop12* and the development of a potentially full-length gene model, representing a major advance over the AaegL3.4 annotation.

### A dramatic increase in the number of known chemosensory receptors

#### Chemosensory receptor overview

Our new genesets of *ORs*, *GRs*, and *IRs* include all previously recognized genes within each family^95–97^. However, we found that 20–30% of previously recognized receptors comprised closely-related pairs that were merged into a single locus in AaegL5 (Fig. 4b and Supplementary Data 20). These were presumably found in regions of the AaegL3 assembly where divergent haplotypes were erroneously represented on separate contigs. We note that the AaegL4 assembly^5^, although not annotated, would likely have resolved many of the same issues. Previous experimental work showed that one pair of similar AaegL3.4 genes (*AaegOr4*, *AaegOr5)* do indeed segregate at a single locus in *Ae. aegypti*^39^.

Six of the new genes and 4 of the new isoforms (private exons only) were missing or fragmented in AaegL3 (Supplementary Data 20). The rest were present but not recognized. One of the two new *OR* genes, *AaegOr133*, has a 1-to-1 orthologue in the malaria mosquito *An. gambiae* (*AgamOr80*, Extended Data Fig. 4), has two relatively close paralogues in *D. melanogaster* (Extended Data Fig. 4), and is one of the most highly expressed *ORs* on female, and particularly male, antennae (Extended Data Fig. 8). Several of the new *IR* genes also fall in clades with clear relatives in *D. melanogaster* (Fig. 4c), and these tend to be expressed in antenna (*AaegIr75l* and *AaegIr31a2*) or proboscis (*AaegIr7p, AaegIr7q, AaegIr7r*) or both (*AaegIr41m*) (Extended Data Fig. 7). Since we identified a significant number of new receptor genes, we decided to revise the naming scheme for *IRs* in both mosquito species. In contrast, we left *OR* and *GR* names intact with the exception of a handful of *GR* isoforms, dropping names for merged genes, and beginning the numbering for new genes where the old genesets left off. Old names for all genes and transcripts are listed in (Supplementary Data 20).

In addition to adding new genes, we also updated the models of a substantial number of previously recognized genes. Most significantly, we extended or added exons to 60% of all previously recognized *IRs* (49 of 81 genes, Supplementary Data 20), resulting in an average protein length increase of over 200 amino acids and greatly narrowing the length distribution for the *IR* family as a whole (data not shown). We also completed the models for 5 *OR* genes that were designated ‘partial’ in AaegL3.4 (ref. ^95^), made major changes to the N-terminus of 8 *GR* genes and 1 *GR* isoform, and made minor changes to the start sites and splice junctions of numerous genes in all three families (Supplementary Data 20). These changes were made manually based on extensive RNA-Seq data^38^ and careful search of flanking sequences for homology to other receptors.

The AaegL5.0 sequences for 63 of a total of 359 receptor transcripts in our new annotation set include loss-of-function mutations that should render them pseudogenes. However, we infer that 20 of these cases likely reflect within-species polymorphism and another 9 result from sequencing/assembly errors. We chose to include updated, intact alleles for these receptors in our genesets (Supplementary Data 20–23) and refer to these sequences as either ‘corrected’ (minor difference between AaegL5 and updated allele) or ‘fixed’ (major difference between AaegL5 and updated allele) (‘C’ and ‘F’ suffixes in Fig. 4c, Extended Data Fig. 4, and Supplementary Data 20). After accounting for these updates, we are left with 34 receptor transcripts that we consider pseudogenes – 11 of 117 *ORs*, 12 of 107 *GRs*, and 11 of 135 *IRs* (‘P’ suffixes in Fig. 4c, Extended Data Fig. 4, and Supplementary Data 20).

#### Large increase of AaegIRs and reannotation of AgamIRs

In light of our recognition of many new *IR* genes in *Ae. aegypti*, we re-examined the *An. gambiae* genome and discovered 64 new *AgamIR* genes to add to the 46 previously described *AgamIRs*^97,103^, bringing the total to 110. Because 6 of these are pseudogenic, the functional *IR* repertoire in *An. gambiae* appears to be 104 proteins. Some of these new genes are related to conserved *IRs* in *D. melanogaster*, for example, *AgamIr87a* (an intronless gene on a 52 kb scaffold in the original PEST strain assembly that was not included in the PEST chromosomal assembly). *AgamIr31a* has a divergent relative immediately downstream of it in chromosome 3R, so the original was renamed *AgamIr31a1*, and the newly recognized gene *AgamIr31a2*. But the vast majority of the new genes, as with *Ae. aegypti*, are related to the genes originally named *AgamIr133–139*, *AgamIr140.1/2*, and *AgamIr142*. In our tree (Fig. 4c) these and the large number of new *Ae. aegypti IRs* are confidently related to the clade of divergent *IRs* in *D. melanogaster* that have been demonstrated to be candidate gustatory receptors and called the *Ir20a* clade (apparently for the lowest numbered IR in the clade)^37,132^. Like the *Ir7* clade, which are also candidate gustatory receptors in *D. melanogaster*^97^, this clade appears to have expanded independently in *D. melanogaster* and mosquitoes. Even comparing these two mosquitoes, multiple expansions of sublineages of the clade have occurred in the anopheline versus culicine lineages, suggesting that gene duplicates have been retained to perceive different chemicals relevant to the chemical ecology of each species. It is noteworthy that all six pseudogenic *AgamIRs*, 9 of the 10 pseudogenic *AaegIRs*, and all 4 of the pseudogenic *DmelIRs* belong to this rapidly-evolving clade, supporting the idea that this clade has undergone rapid gene family evolution, with some receptors being lost to pseudogenization (and likely some were lost from each genome completely, for example, *Ae. aegypti* has lost the relative of *AgamIr105*) when their ligands no longer were relevant to the species’ chemical ecology.

Previously recognized *GR* genes Most of the 114 *GRs* previously described^96^ were present in the new assembly, however 16 *AaegGR* protein names have been dropped as they were nearly identical duplicates of other genes and are not present in the new genome assembly (*AaegGr12P*, *AaegGr24P*, *AaegGr28*, *AaegGr38P*, *AaegGr40a-h*, *AaegGr51*, *AaegGr52P*, *AaegGr70*, and *AaegGr71*). They were all on separately assembled scaffolds, presumably assemblies of alternative haplotypes. The departure of these models disrupts the naming conventions employed earlier^96^. Furthermore, now that the arrangement of these genes on the chromosomes is known, the names are often “chromosomally” and “phylogenetically” jumbled. Nevertheless, this is a problem shared with many arthropod draft genomes, e.g. *An. gambiae*^102^, and even the *D. melanogaster* chemoreceptors, which were named for their cytological locations and hence have some “chromosomal” rationale, are “phylogenetically” jumbled^104^. The original *AaegGR* names have been employed in studies of phylogenetic comparison^22^ and expression^38,133,134^. We therefore chose to retain the original gene numbers, dropping the departed duplicates with higher number and not replacing them, and adding newly recognized genes with the next number in order.

#### Two new *AaegGRs*

Two previously unrecognized divergent *AaegGR* genes were discovered. *AaegGr80* was discovered as an apparently co-transcribed gene with *AaegGr72* (there are just 98 bp between the stop codon of *AaegGr72* and the start codon of *AaegGr80*). This locus was previously modelled in NCBI as LOC110680332. The ancestral final short exon of *GR* genes contains a conserved TYhhhhhQF motif, where h is any hydrophobic amino acid, except in the sugar receptors where the motif is TYEhhhhQF^135^, precisely six codons after a nearly universally present phase-0 intron^129,136^. TBLASTN searches of the genome seeking additional new *GRs* were therefore performed using the amino acids encoded by this final exon from representative *GRs*, with LQ before them to represent the consensus bases of a phase-0 intron 3’ acceptor site (TTGCAG). To increase sensitivity for these searches the default parameters were modified, raising the Expect Threshold from 10 to 1000, reducing the Word Size from 6 to 2, and removing the Low Complexity Filter. These searches revealed one more new gene, *AaegGr81*, discovered with *AaegGr80* as query.

#### Four new *AgamGRs*

There is an unannotated relative of *AaegGr81* in the *An. gambiae* genome, on chromosome 2R from 457,227–458,928 bp, which is a neighbour of a cluster of annotated *GRs*, including *AgamGr58–60*, and so was named *AgamGr61*, the next available number. *Cx. quinquefasciatus* also has a previously unrecognized relative of this gene, here named *CquiGr78*. *An. gambiae* also has an unannotated relative of *AaegGr80*, on chromosome 2R from 54,382,599–54,383,860 bp and immediately downstream of *AgamGr54*, which we name *AgamGr62*, but *Cx. quinquefasciatus* has apparently lost this gene. Furthermore, an unannotated relative of the highly divergent *AaegGr79* was recognized in the *An. gambiae* genome, on chromosome 3R from 44,045,062–44,046,334 and named *AgamGr63* (*Cx. quinquefasciatus* again has no orthologue). It has no *GR* neighbours, and is partially modelled as AGAP028572. Finally, a fourth previously unrecognized *An. gambiae GR* was communicated by Xiaofan Zhou (personal communication), having been discovered (along with independent discovery of *AgamGr61–63*) as part of the 16 *Anopheles* species genome project^80^ and is named *AgamGr64*. It is located on chromosome 3R from 43,704,508–43,705,788bp and is near the *AgamGr9–12* genes (the culicines have no orthologue). These four new *An. gambiae GR* gene models have been communicated to VectorBase to be incorporated in future *An. gambiae* gene sets.

#### Cleaning-up and renumbering alternate

GR isoforms An additional complication to this improvement of the chemoreceptor gene models in the new genome assembly arises in the *GR* family, which has eight alternatively-spliced loci, a phenomenon recognized with the description of the family in *D. melanogaster*^104,136^and present in many other insects including *An. gambiae*^102^ and *Cx. quinquefasciatus*^22^. These isoforms consist of one or more exons encoding the N-terminus of a *GR* spliced to a single set of exons encoding the C-terminus, and the deep RNA-Seq coverage^38^ provides support for most of them. Unfortunately, some of these alternatively-spliced exons were separately assembled in the original assembly, and hence were not associated with the relevant locus, while the large and near identical *AaegGr39/40* loci were duplicates that are now resolved into one locus with eight isoforms. These and other issues require renumbering of the isoforms for several such loci.

#### Updated GR gene models

As described above, *AaegGr80* and *Aaeg81* have been added to the family. Three genes (*AaegGr48*, *AaegGr50*, and *AaegGr75*) are now intact in the new assembly, versus being pseudogenes in the original, and one gene model (*AaegGr45*) is newly recognized as a pseudogene (an intron interrupting the first exon was previously incorrectly modelled to remove a stop codon). Another five gene models were improved (*AaegGr4*, *AaegGr5*, *AaegGr7*, *AaegGr35*, and *AaegGr37*), largely in light of the transcript and RNA-Seq alignments, while the assembly provided two exons that were missing from *AaegGr30* (although those were built from raw genome reads previously). In addition, several proteins resulting from alternative splicing of loci have been modified or added. The *AaegGr39a-h* and *AaegGr40a-h* loci were near identical in the original assembly, but differed in having different isoforms pseudogenized. The new assembly has only one version of this locus, which retains the *AaegGr39a-h* name, with three intact isoforms (a, c, and e) and five pseudogenic ones (b, d, f-h). Finally, Ae. aegypti has a complicated set of alternatively-spliced genes (*AaegGr20a-m*, *AaegGr60a-d*, *AaegGr 61a-c*, and *AaegGr62*) related to the alternatively-spliced *AgamGr37a-f* gene. *AaegGr60–62* were single isoform genes in the original annotation, and while *AaegGr62* remains that way, three additional isoforms are now recognized for *AaegGr60a-d* and two more for *AaegGr61a-c*. Furthermore, the neighbouring *AaegGr20* locus has acquired two more isoforms for a total of 13 (it had isoforms a-k and now has isoforms a-m with the identification of a new isoform after h, now called i, that was so divergent it was not recognized previously, and an ignored pseudogenic fragment before k that is now intact and named l - the other isoforms are renamed to accommodate these).

#### Fixed/corrected GR genes

An additional complication is that for six genes the new assembly does not accurately reflect the genome, as indicated both by comparison with the original assembly, and with a lane of Illumina reads from a single individual and/or available transcript sequences in the TSA^38^. One of these is a base change of an intron 3’ acceptor site from CAG to CAT (*AaegGr17*), and four are single-base frameshifting indels in homopolymers in exons (*AaegGr53*, *AaegGr55*, *AaegGr66*, and *AaegGr72*) (the single individual is heterozygous for most of these mutations). Instead of treating these genes as pseudogenes, their sequences were corrected to encode an intact protein. Another problem is presented by *AaegGr25*, which is intact in the original assembly and the sensory transcriptome, but has suffered an insertion of a 500 bp repeat present widely in the genome, so the intact version is employed herein. A particularly difficult situation is presented by *AaegGr63*, for which the new assembly is seriously compromised by numerous single-base indels (presumably because it is covered by one or very few Pacific Biosciences reads). This gene was therefore modelled based on the original genome assembly. The available transcripts for this gene and its head-to-head neighbour, Gr64, also suggest models that have major length differences from all other *GRs*, so again the original gene models and proteins were employed for them. Finally, while some genes in the new assembly have identical sequences to the old assembly, others have up to several percentage sequence difference, and with the exceptions noted above, the new gene sequences were employed.

#### Summary of *GR* analysis

The final result is that we annotate a total of 72 genes potentially encoding 107 proteins through alternative splicing of 8 loci, but 12 of these are pseudogenic, leaving 95 apparently functional *GRs*. The *An. gambiae GRs* total 93 proteins from 64 genes, none of which are obviously pseudogenic. All of our *AaegGR* proteins, as well as the four new *AgamGRs* and *CquiGr78*, are provided in fasta format in Supplementary Data 22–23.

#### Relationships of *AaegGRs* to *DmelGRs* and *AgamGRs* including biological roles

Phylogenetic analysis of these *AaegGRs* along with those of *An. gambiae* and *D. melanogaster* reveals diverse aspects of the evolution of this gene family in these mosquitoes (Extended Data Fig. 4). While the three carbon dioxide receptors are highly conserved single orthologues in each mosquito^137,138^, there has been considerable evolution of the sugar receptors^135^, including pseudogenization of two genes in *Ae. aegypti* (*AaegGr8* and *AaegGr13*) and loss of a gene lineage from *An. gambiae*. Four other clades of mosquito *GRs* have clear relatives in *D. melanogaster* that likely inform their biological roles. First, *AaegGr34* along with *AgamGr25* are highly conserved orthologues of *DmelGr43a*, a fructose receptor expressed in both peripheral gustatory neurons and within the brain^139^. Second, *AaegGr19a-c* is an alternatively-spliced locus encoding three quite similar proteins with single orthologues in *An. gambiae* (*AgamGr33*) and this lineage is related to *DmelGr28a* and the alternatively-spliced *DmelGr28bA-E*, genes that also have unusual expression patterns beyond peripheral gustatory neurons^140^, and *DmelGr28bD* is involved in temperature sensing in flies^141^. Third, *AaegGr37* and the alternatively-spliced *AaegGr39a-h* locus, along with *AgamGr9a-n*, *AgamGr10-AgamGr12*, and *AgamGr64*, are related to *DmelGr32*, *DmelGr68*, and *DmelGr39aA-D*, proteins implicated in contact pheromone perception in flies and regulation of mating and aggression^142–144^. The complex evolution of these often alternatively-spliced loci mirrors that of the *DmelGr39a* locus within the *Drosophila* genus^145^. Another *D. melanogaster GR* implicated in mating behaviour, *DmelGr33a*^146,147^ has a convincing *An. gambiae* relative in *AgamGr43*, but has been lost from the culicines. Fourth, *AaegGr14* and *AgamGr2* are highly conserved orthologues of *DmelGr66a*, a well-known bitter taste receptor^148^. Most of the remaining *DmelGRs* are implicated in perception of bitter tastants^149–151^, and the same is likely true of many of the remaining mosquito *GRs*, some of which have complicated relationships with *DmelGR* lineages, for example, these mosquitoes each have three *GRs* (*AaegGr16–18* and *AgamGr7*) that cluster with *DmelGr8a*, which participates in perception of a plant-derived insecticide, L-canavanine^148^. Surprisingly, *Ae. aegypti* and *Cx. quinquefasciatus* have lost the orthologue of *AgamGr43*, which is apparently related to *DmelGr33a* but does not share microsynteny with it, a well-known bitter taste receptor that is also involved in courtship behaviour ^146,147^.

The remaining relationships of these mosquito *GRs* are typical of insect chemoreceptors (Extended Data Fig. 4), ranging from highly conserved single orthologues comparable to the carbon dioxide or fructose receptors (e.g. *AaegGr73*/*AgamGr53* or *AaegGr30*/*AgamGr47*) whose ligands and biological roles are likely to be shared across these mosquitoes but which were apparently lost from drosophilids, to instances of loss from one or more lineages (e.g. the *Ae. aegypti* orthologues of *AgamGr1*, *AgamGr34*, *AgamGr58*, *AgamGr59*, and *AgamGr60* were lost), to major gene-lineage-specific expansions in each species. The three most prominent of the latter are the independent expansions of *AgamGr55*/*AaegGr74a-e* and *AgamGr56a-f*/*AaegGr67a-e/ AaegGr68/ AaegGr69*, an expansion of 15 *AegGRs* related to *AgamGr40*, and the clade that includes *AaegGr20a-m*, *AaegGr60a-d*, *AaegGr61a-c*, and *AaegGr62*, all of which are neighbours in chromosome 3, and related to *AgamGr37a-f*. The latter expansion of these proteins from 6 in *An. gambiae* to 21 in *Ae. aegypti*, and an even larger number in *Cx. quinquefasciatus*^22^, surely indicates an important involvement in idiosyncratic aspects of the chemical ecology of culicine mosquitoes.

